# Unraveling lncRNA Diversity at a Single Cell Resolution and in a Spatial Context across Different Cancer Types

**DOI:** 10.1101/2024.08.12.607523

**Authors:** P. Prakrithi, Tuan Vo, Hani Vu, Zherui Xiong, Loan Nguyen, Andrew Newman, Vicki Whitehall, Jazmina L. Gonzalez Cruz, Ishaan Gupta, Quan Nguyen

## Abstract

Long non-coding RNAs (lncRNAs) play pivotal roles in gene regulation and disease, including cancer. Overcoming the limitations of lncRNA analysis with bulk data, we analyzed single-cell and spatial transcriptomics data to uncover 354937 novel lncRNAs and their functions across 13 cancer types. LncRNA functions were assessed by identifying their cell-type specificity and distinct spatial distributions across different tissue regions. First, lncRNAs were computationally validated by comparing to existing databases, and experimentally validated using spatial long read sequencing methods. Further, genome-wide computation of spatial-autocorrelation identified coexpression of lncRNAs with cancer-associated protein coding genes across the tissue. Additionally, genomic co-localization of lncRNAs with regulatory features and disease-associated genetic variants suggest possible functional association. The identified lncRNAs were analyzed for responses to immunotherapy and prognostic value, revealing cancer-outcome associated lncRNAs. We have made this novel resource available as an open website ‘SPanC-Lnc’ hosted on AWS cloud to serve as a pan-cancer atlas of single cell- and spatially-resolved lncRNAs. These can complement established biomarkers because they reflect the unique characteristics of specific cell populations within tumors, offering new insights into disease progression and treatment response.

---

Long non-coding RNAs (lncRNAs) constitute the vast majority of the permissively transcribed genome. Despite this, the majority of our knowledge about transcriptional events is limited to the 1-2% of the genome that encodes proteins^1^. With advancements in high throughput technologies and several key studies, it has become increasingly apparent that lncRNAs are of functional relevance, contributing to diseases like cancer where they are often misregulated^2^.

LncRNAs are involved in several functions at epigenetic levels including DNA methylation, histone modification and chromatin remodeli ^3^. These mechanisms influence and control the expression of certain factors regulating disease progression such as genes pivotal to DNA repair, apoptosis, autophagy, transcription, cell-cycle regulators and signaling pathwa ^3^.

LncRNAs can also hybridize with pre-mRNAs, blocking the recognition of splice sites by spliceosomes, thereby modulating their alternative splicing to produce alternate transcripts^4^. Cytoplasmic lncRNAs typically function as microRNA (miRNA) sponges, modulating the expression levels of nearby miRNAs^4^.

Moreover, lncRNAs can bind to proteins, thereby affecting their localization, activity and protein-protein interaction^5^. For example, lncRNA KILR sequesters RPA1 and inhibits its movement to sites of double strand breaks^6^. Despite being mostly non-coding, putative small open reading frames in a subset of lncRNAs can be translated into a polypeptide. For instance, Wang et al.^7^ have identified that LINC00908 encodes a differentially expressed polypeptide in triple-negative breast cancer (TNBC), named as the endogenously expressed polypeptide ASRPS. ASRPS directly binds to the coiled-coil domain (CCD) of STAT3 thus inhibiting STAT3 phosphorylation, leading to a decrease in expression of the vascular endothelial growth factor (VEGF) and inhibition of tumor angiogenesis in breast cancer ^7,8^.

LncRNAs can play dual roles as either oncogenes or as tumor suppressors. For example, *HOTAIR* is involved in tumorigenicity in pancreatic cancer and can also cause proliferation and metastasis in colorectal cancer^9^. It is also associated with poor prognosis in several cancers. The lncRNA *PCA3* is the only FDA-approved lncRNA biomarker and is used in diagnosing prostate cancers^10,11^ (as of October 2023). LncRNAs like *NORAD* and *PANDA* can suppress transcription factors, thereby inhibiting the expression of targeted genes. *NORAD*, for instance, binds and chelates the calcium-binding protein S100P, thus inhibiting its metastasis-promoting signaling network^12^. Conversely, *PANDA* is involved in the DNA-damage response by interacting with the nuclear transcription factor Y subunit A (NF-YA) potentially preventing it from activating apoptotic gene expression^13^. Similar mechanisms have also been observed for other lncRNAs^6^.

LncRNAs serve as a valuable class of molecular targets for disease identification. However, their detection has been constrained by factors such as their low abundance, cell-type-specific expression patterns, and the reliance of existing computational tools on prior annotations^14^. Previous studies on lncRNAs have predominantly relied on bulk RNA-seq data, which unfortunately results in the loss of cell-type information and tissue spatial context during sample preparation. However, understanding the spatial context of gene or lncRNA expression is important for elucidating the transcriptional regulation in both developmental and diseased states.

It provides valuable insights into a transcript’s location within a tissue, its neighboring cells, colocalizing transcripts or proteins and their interacting partners, thereby enhancing our comprehension of biological processes. Hence, bulk RNA-seq imposes limitations on our ability to fully delineate the functional consequences of each lncRNA.

Thanks to advanced sequencing technologies such as single-cell sequencing (scRNA-seq) and spatial transcriptomics (ST), researchers can now investigate transcript expression at the single-cell level while preserving spatial information. The 10x Visium technology allows the capture of the polyA tails of transcripts. Up to 40% of lncRNAs are polyadenylated and an additional 40% of them are bimorphic^15^. Thus, using this protocol, such transcripts could be captured and analyzed. Recent studies using scRNA-seq for breast cancer samples 16,17 have evaluated the annotated lncRNAs at a single-cell level. Other studies claiming to have analyzed the ‘spatial transcriptome’ of lncRNAs have typically examined lncRNA expression across different bulk tissues, rather than capturing the nuanced expression patterns of lncRNAs within a tissue section^18,19^. Unfortunately, these studies also lack the capacity to explore novel unannotated lncRNAs. Several methods for novel transcript identification have been established^20,21^. Applying these methods to ST and scRNA-seq data would not only facilitate the expansion of annotations but also unveil additional layers of information regarding lncRNAs, including cell-type specificity and spatial context. Although these technologies have not yet addressed the identification of non-polyadenylated lncRNAs or the entire spectrum of rare lncRNAs, creating a repository of potential lncRNAs detected through cutting-edge spatial transcriptomics technologies would establish a robust foundation. Such a resource would serve as a pivotal reference point for further investigation and development within the scientific community.

In this study, we combined large datasets from the recent spatially-resolved RNA sequencing modality 10x Visum and single-cell resolution scRNA-seq by 10x Chromium to discover lncRNAs in tissues from 13 different cancer types. We demonstrate new types of analyses that can be performed on such data to identify the potential functions of lncRNAs. While previous pan-cancer studies have examined clinically relevant lncRNAs^22–24^, this study represents the first effort to analyze novel potential lncRNAs incorporating spatial context and cell-type information in cancer research. The annotations and lncRNA expression data of each tissue have been made accessible through the interactive website **‘SPanC-Lnc’**: <u>**S**</u>patial and <u>**S**</u>ingle Cell <u>**Pan**</u>-<u>**C**</u>ancer Atlas of <u>**lnc**</u>RNAs. Researchers and clinicians can leverage these resources to gain deeper insights into the roles of lncRNAs in cancer biology and to identify potential biomarkers and therapeutic targets.

## RESULTS

### Identification of potential lncRNAs in spatial and single cell datasets

Samples from 36 in-house ST (new or previously published) and scRNA-seq datasets across 13 cancer types were analyzed for potential novel lncRNAs (**Fig. 1a)**. The cumulative expression of all unannotated Transcriptionally Active Regions (uTARs) detected in each sample was projected on the corresponding tissues and compared with that of the cumulative coding gene expression. Three representative examples for different cancer types are displayed in **Fig. 1b**. uTAR and coding gene expression for the remaining samples are also shown in **Supplementary Fig. 1 and 2** respectively. The expression of several previously identified cancer-specific lncRNAs was also visualized **(Supplementary Fig. 3)**. It is worth noting that while lncRNAs like *MALAT1* and *NORAD* were abundantly expressed across the tissue sections in various samples **(Supplementary Fig. 3a)**, others displayed a higher degree of tissue specificity. For instance, both *PICSAR* and *LINC00520*, known to be squamous cell carcinomas-specific^25^, were markedly more expressed in cutaneous Squamous Cell Carcinoma (SCC) and Head and Neck Oropharyngeal SCCs (H&N OPSCC) compared to the other cancer types. In contrast, while *PICSAR* and *LINC00520* were either absent or expressed at lower levels in breast and colorectal cancer, *TINCR* (breast cancer specific^26,27^) was highly expressed in the breast cancer samples, **(Supplementary Fig. 3b)**. Similarly, *KCNQ1OT* (colorectal cancer-specific^28,29^) showed higher expression levels in colorectal cancer samples compared to the lncRNAs specific to the other cancer types. The datasets used and the number of uTARs identified are summarized in **Fig. 1c**. The numbers vary depending on the cancer type and coverage. For example, a higher number of uTARs were found in the breast cancer (∼45,000-60,000), one of the kidney cancer samples (namely KidA) (∼45,000) and in the NSCLC samples (162,588, not displayed in figure due to high coverage and hence very high uTARs detected) while around 15,000-25,000 were detected in the other cancer samples. Overall, more uTARs were detected with scRNA-seq compared to ST, as scRNAseq generally captures a higher number of cells. In most cases, approximately 40-60% of the identified uTARs overlapped with lncRNAs cataloged in two selected lncRNA public databases such as FANTOM, LncExpDB (**Fig. 1c**) and NONCODE (**Supplementary Fig. 4**). The remaining transcripts were likely novel discoveries (**Fig. 1c**). These results suggest the reliability of our lncRNA detection method utilizing spatial data and suggest that the novel uTARs identified merit further investigation. 210 uTARs were expressed on a pan-cancer level (i.e., expressed in at least one sample per cancer type used in the study), while others exhibited specificity to particular cancer types (**Supplementary Fig. 5a, 5b**). The relative uTAR-gene counts for each tissue at single Visium spot-level resolution was calculated as the total number of uTARs divided by the total number of genes detected per spot **(Fig. 1d)**. A higher value indicates a higher number of total uTARs detected per Visium spot with respect to the number of genes. We observed a higher mean uTAR-gene count ratio value in the Melanoma samples, indicating a higher proportion of uTARs per spot relative to coding genes **(Fig. 1d)**. However, this could be due to the lower sample quality (i.e. < 200 median genes per spot). The consistency of detection across different sequencing platforms (Spatial Transcriptomics, 3’ and 5’ scRNA-seq) for Melanoma samples was also analyzed. About 1,205 uTARs were detected in melanoma samples across all the three different platforms **(Supplementary Fig. 5c)**. Since the uTARs analyzed in this study span various cancer types and tissues, we designate them with a nomenclature of “cuTAR” (cancer-associated uTAR), followed by a unique numerical identifier as used in the SPanC-Lnc database. We then investigated some of these potential disease-associated candidates.

**Fig. 1:**
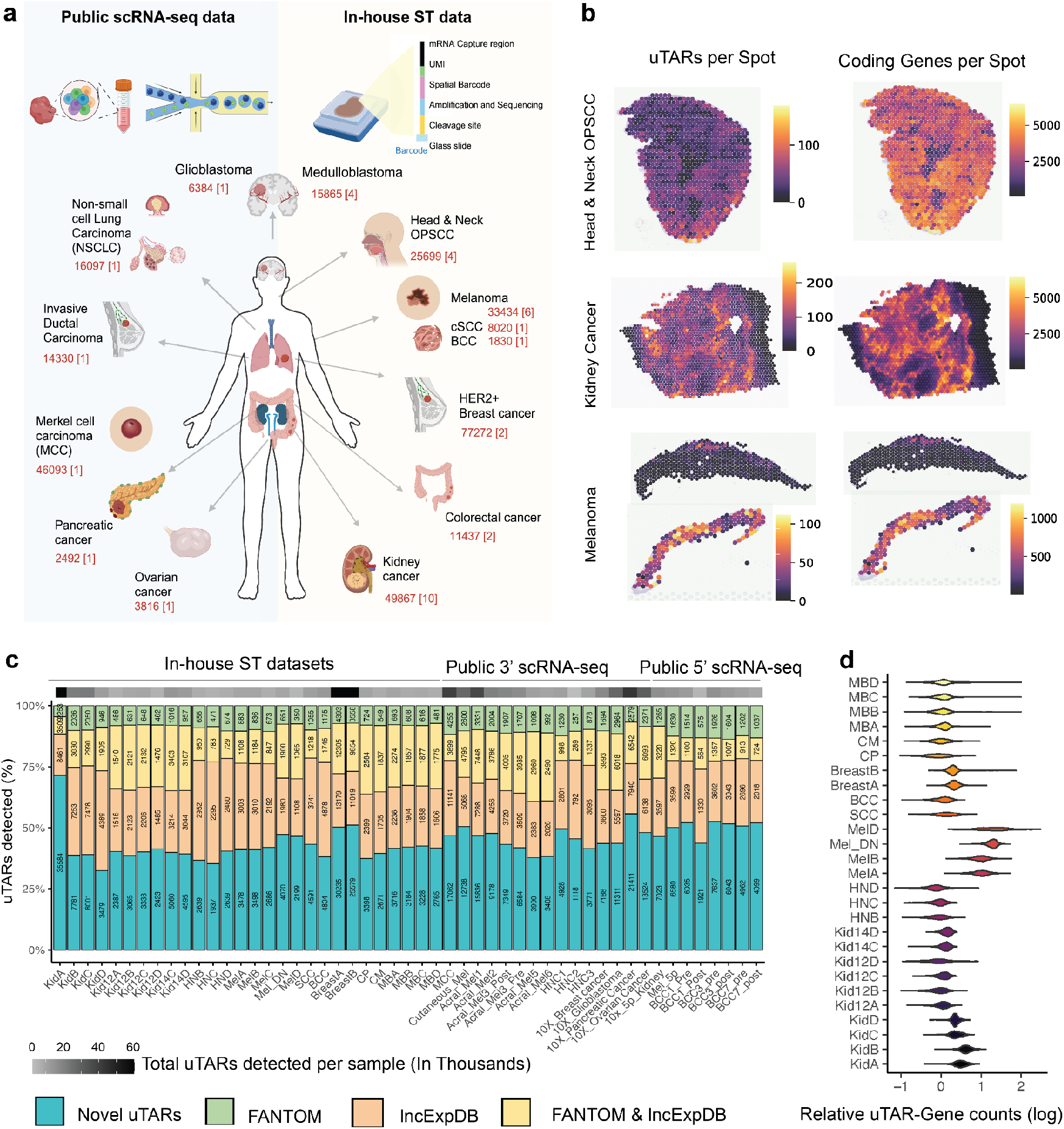
Pan-cancer identification of novel lncRNAs from spatial and single-cell data. **a**, Different cancer types used from public scRNA-seq and in-house spatial transcriptomics datasets. Numbers highlighted in red indicate the number of unannotated Transcriptionally Active Regions (uTARs) detected for each cancer type, followed by the number of samples analyzed within square brackets. **b**, uTAR counts overlaid on the tissue for representative samples as compared to the protein-coding gene counts. **c**, Breakdown of known and novel uTARs identified per cancer sample. The blue region indicates novel uTARs and green, orange and yellow indicate an overlap with public datasets (FANTOM, LncExpDB and both respectively). Bar labels and the grey scale bar on top show the number of uTARs found from each source. **d**, Gene-uTAR expression ratio, (i.e., number of genes vs. number of uTARs), per spot across the ST samples. A higher value indicates a higher number of uTARs detected per spot with respect to the number of genes.

**Fig. 2:**
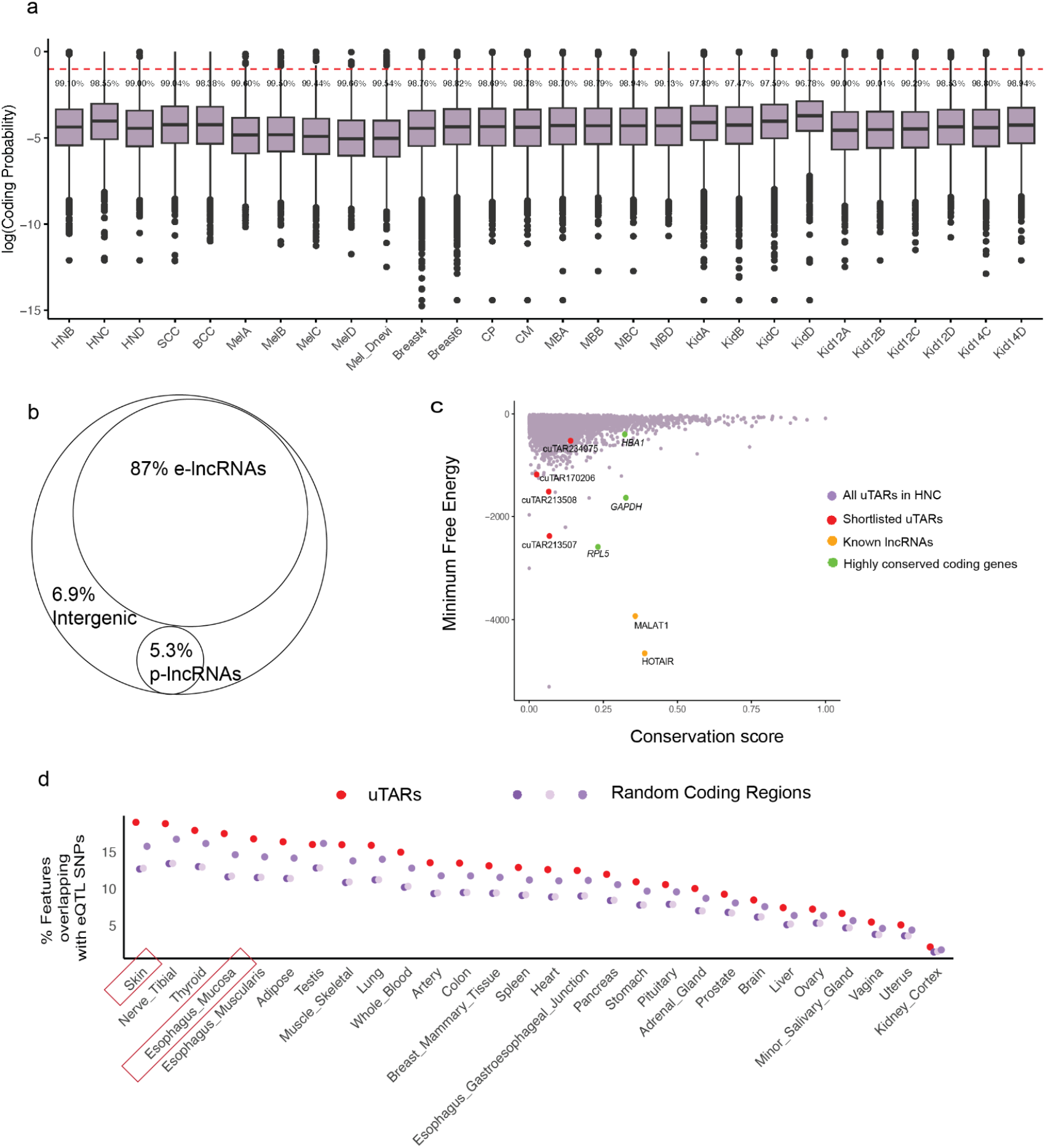
Analysis of sequence features and co-localization with functional SNPs. **a**, Analysis of coding potential of the identified uTARs. Box plots display log coding probability values for each cancer. The red dashed line indicates the log coding probability cut-off. Percentage values show the proportion of non-coding transcripts based on this cut-off. Sequences with a predicted coding potential below the standard cut-off for humans, 0.364,were determined to be non-coding. **b**, Classification of lncRNAs based on their overlap with regulatory features as enhancer-associated lncRNAs (e-lncRNAs), promoter-associated lncRNAs (p-lncRNAs) and intergenic lncRNAs **c**, Conservation and Minimum Free Energy (MFE) calculations as a measure of stability. **d**, Overlap of uTARs and randomly chosen regions from coding genes with tissue-specific eQTLs.

**Fig. 3:**
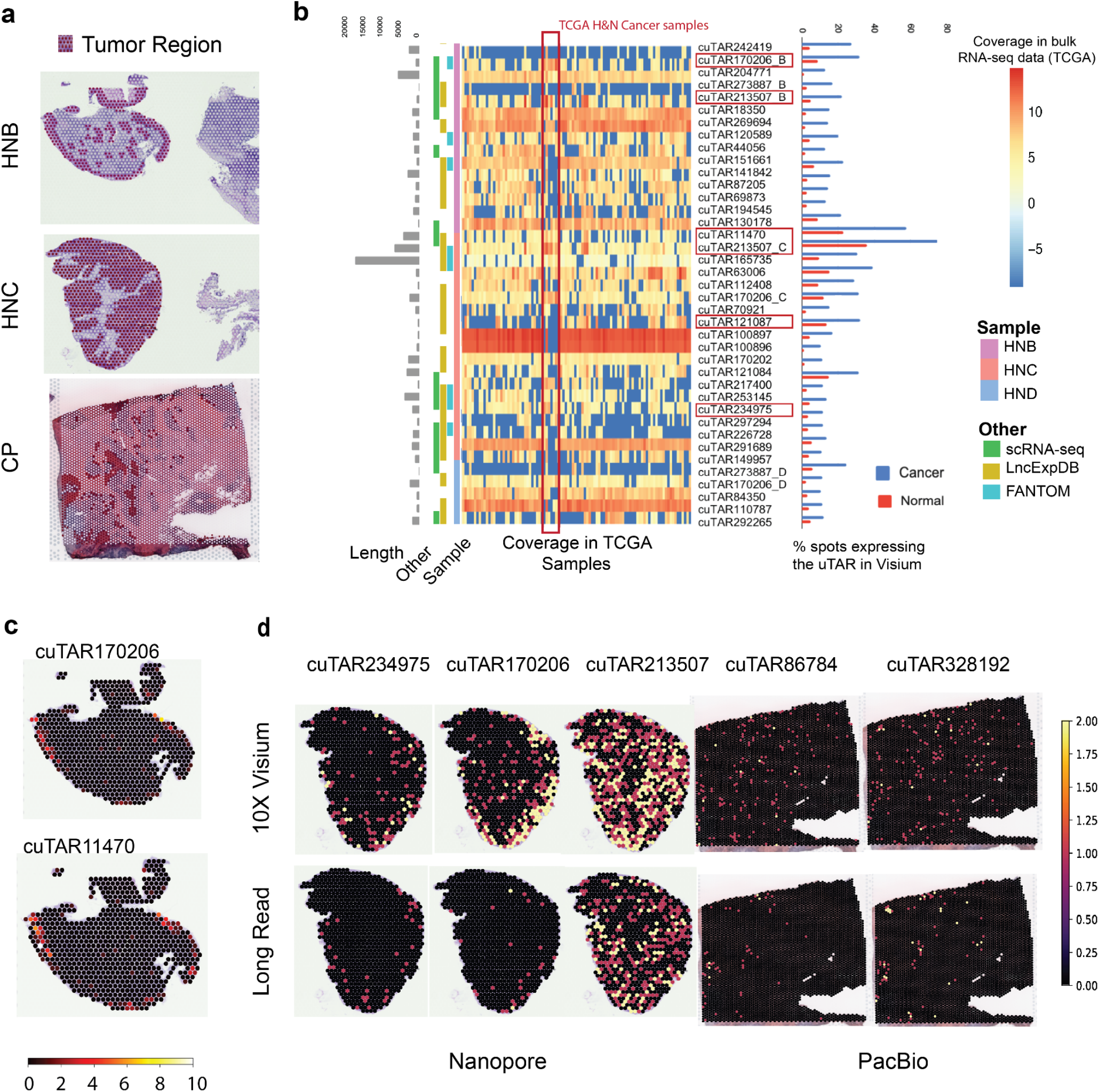
Identification of tumor region specific uTARs and validation with long read sequencing technologies. **a**, Tumor regions (dark shaded) in the Head and Neck cancer samples B (top) and Colorectal cancer Primary tumor CP (middle). **b**, (From left to right) uTAR lengths, overlap of uTARs with that from scRNA-seq data and with lncRNAs from public databases, expression of uTARs in bulk RNA-seq samples from TCGA (normalized bigWig coverage), and quantification of the uTARs in the cancerous and normal regions of Head and Neck cancer samples. The overlapping uTARs across samples are indicated by the sample suffix (_B, _C and _D) The highlighted uTARs with red boxes indicate the ones with higher expression in the tumor region than the normal region, some of which are shown in panel **c**. These show relatively higher expression in the analyzed H&N cancer samples from TCGA highlighted using the vertical red box **c**, Expression of exemplar cancer region-specific uTARs projected on the tissues. **d**, Validation of some uTARs with long read sequencing ONT and SMART-Seq for Head and Neck and Colorectal cancer samples, respectively.

**Fig. 4:**
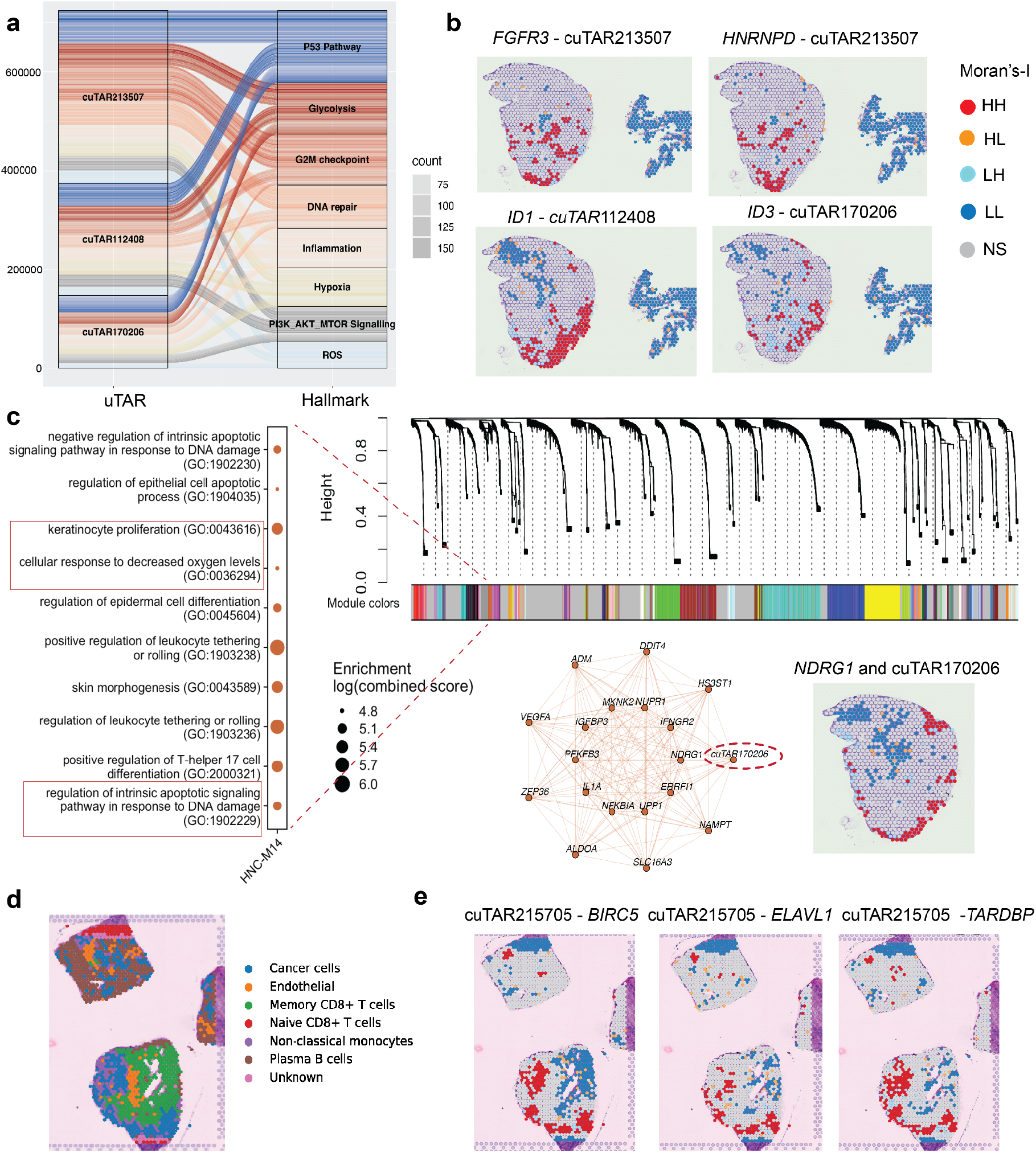
Spatial co-expression of uTARs with cancer relevant genes. **a**, Top uTARs in Head and neck cancer (sample C) with high spatial autocorrelation (HH: both features expressed in a given spot) with different cancer-relevant gene sets. **b**, Spatial correlation of expression with genes (LH/HL: Either the gene or cuTAR is expressed, LL: Both features not expressed, NS: No significant autocorrelation). **c**, WGCNA analysis shows 32 genes displaying high coexpression with cuTAR170206 (circled in red) that forms part of the same regulatory module which includes genes like NDRG1, the downregulation of which is associated with metastasis in OPSCC and other genes involved in DNA repair, hypoxia response and negative regulation of apoptosis (associated GO terms highlighted in red boxes). **d**, Cell-type annotations of a breast cancer tissue. **e**, Co-expression of cuTAR215705 with the mRNA encoding the RNA-binding protein ELAVL1 and its interacting partner *BIRC5* and the mRNA encoding TARDBP5 in breast cancer.

**Fig. 5:**
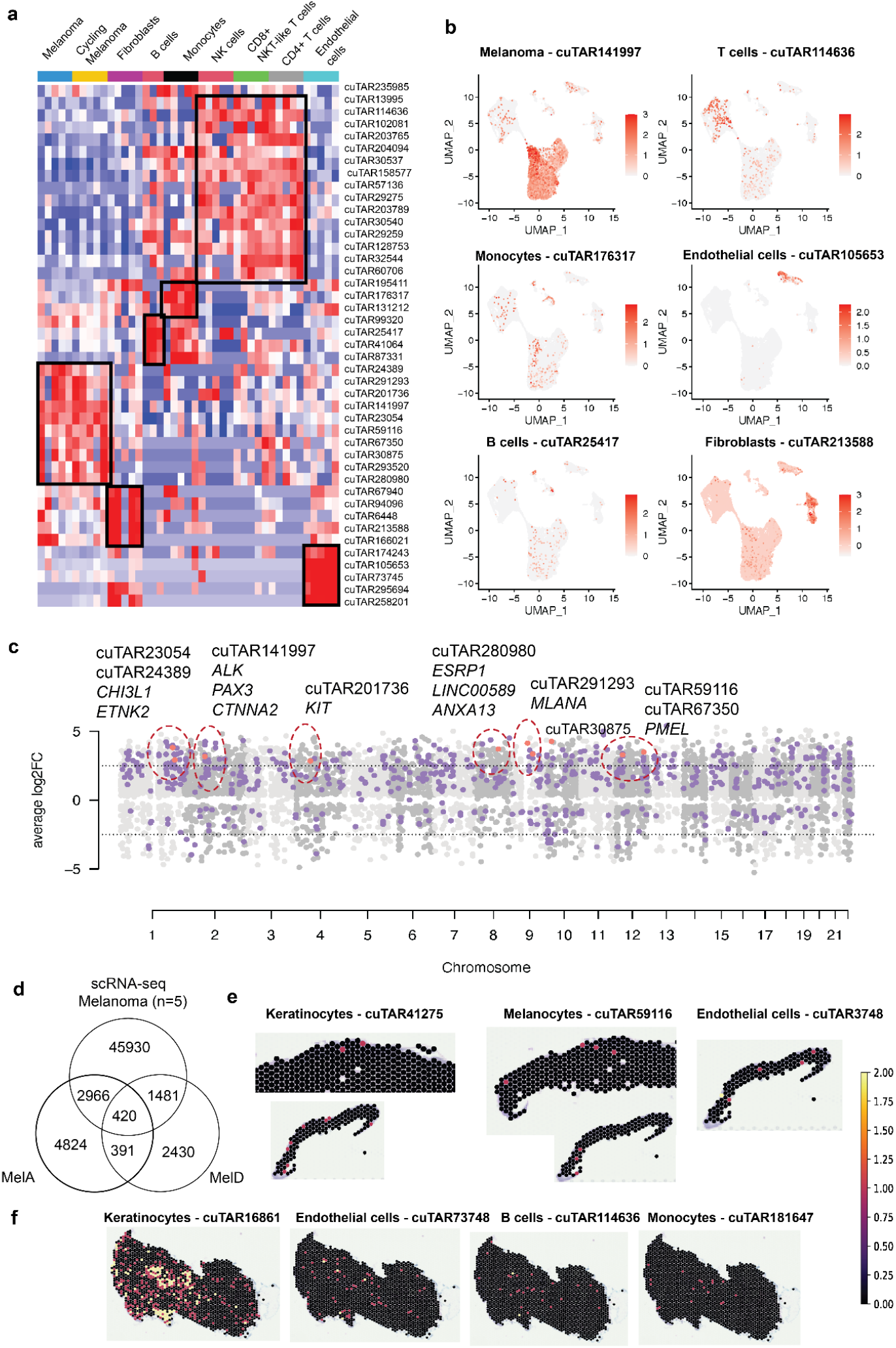
Cell-type specific expression of lncRNAs. **a**, Cell-type specific uTARs (highlighted with black boxes) in scRNA-seq Acral Melanoma samples. **b**, Expression of some cell-type specific uTARs projected on the UMAP. **c**, Expression trends as compared to coding genes. Tumor-specific uTARs and proximal cancer-associated coding genes are highlighted in red circles. **d**, Cell-type specific lncRNAs identified in the scRNA-seq data that overlap with in-house ST data. **e, f**, Expression of cell-type specific uTARs in 10X Visium Melanoma samples (Samples MelA and MelD respectively).

### Classification and transcriptome-wide analysis of functional implications

Analysis of the sequences confirmed that 95.7% of the total uTARs lack coding potential. A sample level distribution is shown in **Fig. 2a**. Moreover, 92.1% of them were longer than 200 bp, suggesting that the majority of uTARs are most likely to be ‘long’ ncRNAs^30^. The lncRNAs with a coding potential might indicate their role in encoding functional micropeptides, as opposed to necessarily being protein-coding transcripts^31,32^. Since the uTARs are only estimated transcript boundaries using data captured from either the 3’/5’ ends, the calculated coding potentials may not be as accurate as using the full-length sequence, but could be still be reflective of their coding potential. Based on their overlap with enhancers from the EnhancerAtlas^33^ and TSS data from FANTOM, the uTARs were classified as enhancer-associated (e-lncRNAs) and promoter-associated (p-lncRNAs). The majority were e-lncRNAs (87%), followed by intergenic lncRNAs (6.9%) and p-lncRNAs (5.3%) (**Fig. 2b**). When a lncRNA overlaps with such cis-regulatory elements, it suggests that the lncRNA may be involved in regulating its associated gene/transcript. Various metrics can help infer the functional significance of a sequence. One such metric is the conservation score, where highly evolutionarily conserved sequences tend to be associated with common essential functions across organisms, while less conserved sequences may exhibit specific functions in an organism. Another metric is the minimum free energy (MFE), which reflects the potential functionality of transcripts. Lower MFE values suggest transcripts with more stable secondary structures, making them more likely to be functional. The conservation and MFE of the uTAR sequences for H&N sample C (HNC) were calculated. Most top uTAR candidates have conservation scores lower than a housekeeping gene *GAPDH* and a known lncRNA *HOTAIR* **(Fig. 2c)**, while cuTAR100897 had a similar conservation score as the latter **(Fig. 2c)**. However, this uTAR had sequences multimapping with other regions of the genome, which could possibly explain the high conservation score and must be carefully evaluated for downstream analysis.

Gene expression can be influenced by genetic variants, with tens of thousands of variants identified to date that are associated with altered gene expression across tissues or cell-types^34^. The overlap of these variants, such as Expression Quantitative Trait Loci (eQTLs) or those identified in genome-wide association studies (GWAS), with non-coding transcripts could suggest potential tissue-specific or disease-associated gene regulation of these lncRNAs. Analysis of uTARs overlapping with eQTLs from the GTEx project^35^ showed that the majority of the lncRNAs from the in-house ST datasets, mostly comprising cutaneous and OPSCC datasets, were enriched for eQTLs that are involved in gene expression regulation in skin (20% of the uTARs) and esophageal mucosa (19%), which is the closest tissue to the oropharyngeal mucosa, as compared to randomly sampled coding regions (highlighted in red boxes) (**Fig. 2d**). Three sets of an equal number of random coding regions were considered and the overlap of eQTLs were calculated. There was a significant lower overlap with three sets of randomly chosen coding regions (Adjusted *p*-value = 7.85e^-29^ from Fisher’s exact test) **(Fig. 2d)**. The eQTLs overlapped with both the pan-cancer uTARs and the cancer-specific uTARs were identified using window sizes of 10kb, 50kb and 100kb. The genes associated with these colocalized regions were then queried in enrichR for enriched pathways, GO terms, cell-types and tissues which are computed using Fisher exact test^36^. It was observed that for the uTARs specific to given cancers, the eQTL-associated genes were enriched in their respective GTEx tissues and cell-types. For example, uTARs specific to brain tumors were enriched for eQTLs associated with the “Brain-Cerebellum Female” category (**Supplementary Fig. 6)**. For the pan-cancer uTARs, while a 10kb window of co-localized eQTLs did not yield many cancer-associated terms, a 100kb window size revealed pathways involved in several cancers **(Supplementary Fig.7)**. Additionally, approximately 10-15% uTARs overlapped with cancer associated GWAS SNPs **(Supplementary Fig. 8)**, suggesting the potential value of further detailed analysis. These findings collectively point towards genetic-level regulation.

**Fig. 6:**
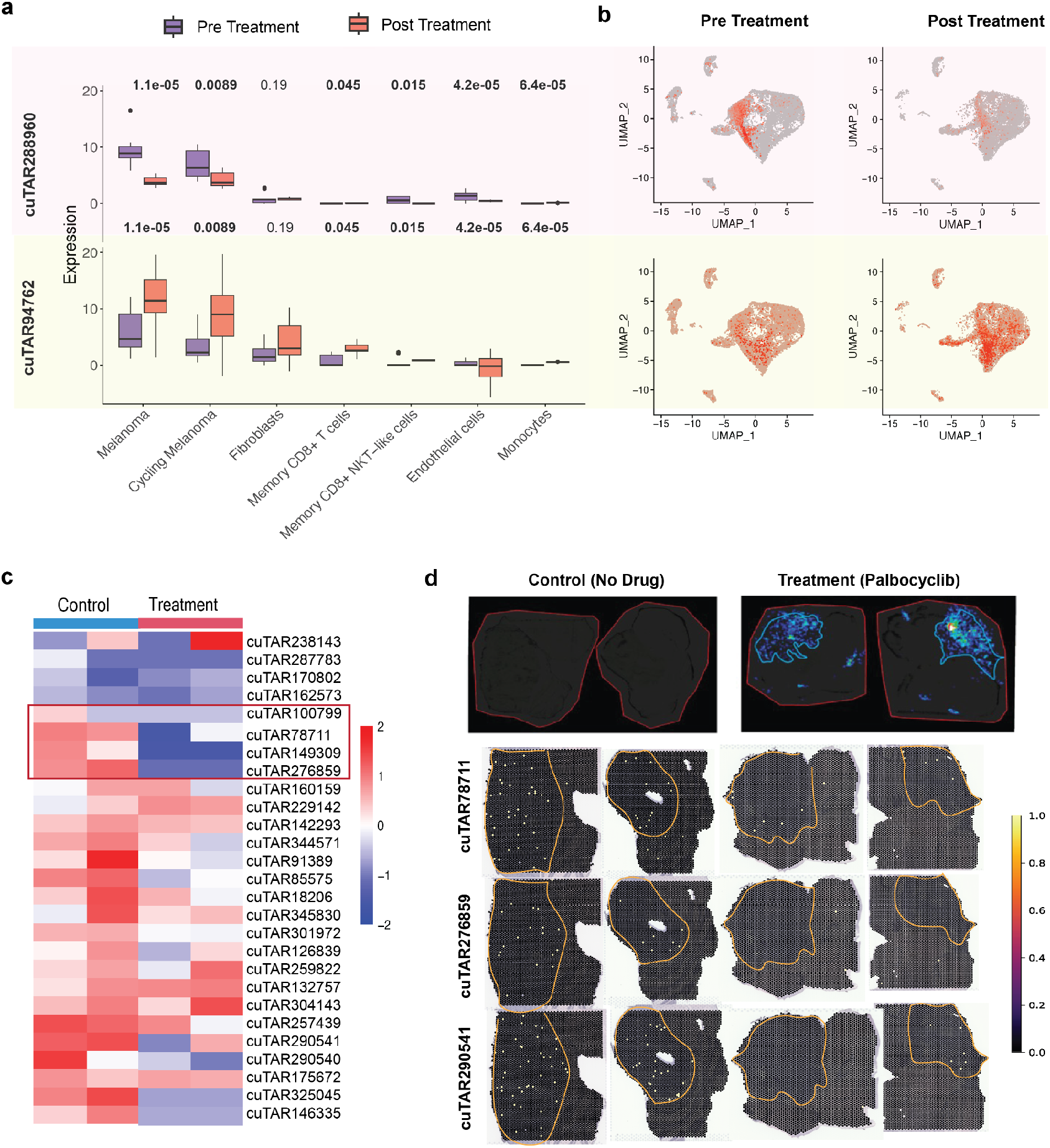
Response of lncRNAs to drug treatments. **a**, Expression trends of melanoma-specific uTARs in response to anti-PD1 therapy **b**, and their UMAP projections. **c**, Downregulation of uTARs in Medulloblastoma in response to Palbocyclib. Some uTARs showing explicit differences in expression are highlighted with the red box. **d**, Spatial expression of the tumor specific uTARs downregulated post treatment highlighted with a red box in **c**. Red outlines define the tissue borders and yellow outlines highlight the human tumor region.

### Identification of lncRNAs enriched in cancer regions within the tissue

Next we asked if the uTAR expression was restricted specifically to the tumor area within the biopsies. The cancerous regions of each tissue were identified from the histological annotations by a pathologist **(Fig. 3a)**. Top cuTAR candidates for each cancer type were shortlisted based on their detection across various datasets and higher expression in the annotated cancerous regions. The percentage of spots in the cancerous *v*.*s*. Normal regions expressing the selected uTARs were calculated **(Fig. 3b)**. Expression of these top cuTAR candidates was detected across many cancer types from The Cancer Genome Atlas (TCGA)^37^ bulk RNA-seq samples. Some uTARs showed higher expression in the two OPSCC samples used, consistent with the observation that all these candidates are tumor-specific in the H&N ST data **(Fig. 3b)**. Among the uTARs differentially expressed across these two annotated clusters **(Supplementary Fig. 9)**, visual examination of gene expression overlaid on the tissues reveal some uTARs with higher expression in the tumor regions as (shown in **Fig. 3a and Fig. 3c)**. Not only cancer region specificity, but also sub-clone specificity was observed. The tumor sub-clones were identified using CopyKAT^38^ **(Supplementary Fig. 10)**.

### Confirmation of lncRNAs using long-read sequencing for spatial transcriptomics samples

Since the 10X Visium protocol captures only the 3’ ends of the fragmented transcripts, the actual transcripts are likely longer than the cuTAR boundaries detected. This gap was addressed by employing Oxford Nanopore Technology (ONT) long read sequencing methods and HiFi long read sequencing with PacBio’s SMRT sequencing to capture full length sequences of lncRNA transcripts present in 10X Visium libraries and corroborate the uTAR signals detected by the standard 10X Visium sequencing. We found the signals detected to be consistent across different technologies, suggesting that these lncRNAs are likely to be genuine signals rather than artifacts from the TAR-Seq pipeline.

Samples from OPSCC, SCC, BCC human cancer Visium samples and PDX mouse Visium samples were used for the ONT experiment. Specifically, from OPSCC (HNC), we obtained 20 million ONT reads, with 3422 spatial barcodes identified by our customized scNanoGPS pipeline, allowing us to map to 1029 Visium spots across the tissue. We successfully recovered 87% of the 20 million reads, identifying approximately 960 lncRNAs present in at least three spatial spots overlapping the tissue. This accounts for 31.7% of those detected using the 10X Visium platform (**Supplementary Table 1**), thus validating the results obtained from the 10X Illumina data. Similar spatial patterns were observed **(Fig. 3d)**, with up to 38% of spots showing expression in both the technologies and up to 86% for coding genes. Interestingly, 166 novel uTARs that were not detected with the short-read data were identified by applying the TAR-Seq pipeline directly on the Nanopore BAM files in addition to the signals confirmed earlier by using the TAR annotations generated using the 10X Visium data. The consistency of expression patterns for coding genes were also checked **(Supplementary Fig. 11)**. There was a significant overlap of expressing spots across both the platforms for some uTARs **(Supplementary Fig. 12a)**. The same analysis was performed for cutaneous SCC, BCC (Basal cell carcinoma) **(Supplementary Fig. 12 b, c)** and medulloblastoma samples. This experiment not only validated the computationally identified lncRNA signals but also could be useful to analyze lncRNA isoforms and their spatial context in the future analyses.

**Table 1:** Comparison of available resources for lncRNAs.

| Resource | FANTOM CAT | lncExpDB | NONCODE v6 | SPanC-Lnc<br>(This study) |
| --- | --- | --- | --- | --- |
| No.of lncRNAs reported | 199375 | 101293 | 173112 | 353141 |
| Source | Primary cell lines and tissues | Primary cell lines and tissues | lncRNA databases including Ensembl, RefSeq, lncRNAdb, LNCipedia, CANTATAdb, GREENC, old versions of NONCODE and literature | Tissues |
| Condition |  | Healthy and disease | Human cancers and plant morphology | 13 cancer types |
| Technology | RNA-seq | RNA-seq | RNA-seq, scRNA-seq | scRNA-seq and Spatial Transcriptomics (10X Visium, Nanopore, PacBio) |
| Organism | Human | Human | Animals and Plants | Human |
| Citation | Hon *et al.* <sup>56</sup> | Li *et al.* <sup>57</sup> | Lianhe *et al.* <sup>58</sup> | This study |

For the HiFi long read sequencing with PacBio’s SMRT sequencing run with cDNA of colorectal cancer samples from the 10X Visium experiment, we generated approximately one million reads. By aligning the reads and tagging them with spatial barcode information, we found up to 1500 lncRNAs per sample, which accounted for 17% of those detected with 10X Visium **(Supplementary Fig. 13)** thereby confirming the findings of the 10X Visium Illumina data. We then examined specific cuTARs and found up to 30% of the spots showed significant consistency in expression for the same set of spots across both the platforms for two colorectal cancer specific uTARs **(Fig. 3d)** (cuTAR86784: 19%, p-value: 0.011; cuTAR114310: 29%, p-value: 0.014). Overall, we found the HiFi yield was better than that of ONT considering the number of reads captured and processed.

### Experimental detection of lncRNA using quantitative reverse transcription PCR

We further experimentally validated and quantified the cuTAR models that were based on short and long read sequencing platforms. With specific primers targeting each of the selected uTARs, we found cuTARs in the H&N-C and -B (HNC and HNB) samples, while a few other cuTARs were tested for the other SCC, BCC and colorectal cancer samples **(Supplementary Fig. 14)**. All the lncRNAs were detected except for cuTAR121087 whose detection seemed to be very low although high expression was detected with Visium for the HNC sample **(Supplementary Fig. 14)**. Both the primers designed for just cuTAR100897 showed off-target regions on UCSC-BLAT and *In-silico* PCR which could attribute to the higher expression detected, for with ONT sequencing also showed many multi-mapping reads and hence was excluded in the analysis. All these experiments help accurately identify high confidence lncRNAs. Considering gene expression levels, we found a generally consistent trend observed with the 10X Visium data **(Supplementary Fig. 14)**.

### Inferring potential functions by spatial co-expression analysis of lncRNAs with cancer-associated genes

Transcripts that are expressed together tend to function together. To identify the cuTAR-gene pairs co-expressed in space, we first identified spatially-variable uTARs, those that were most likely to change with pathological heterogeneity across the tissue (**Supplementary Fig. 16)**. For data-driven detection of functional uTARs, we used Bivariate Moran’s Index to screen for gene-pairs between the spatially-variable uTARs and cancer-hallmark genes. Our spatial co-expression approach suggests possible co-regulation gene networks between unknown uTARs and known cancer-associated genes. The top three uTARs with more than 50 high-high (HH) spots showing co-expression with a greater number of cancer-relevant genes compared to the other uTARs in the H&N sample C (HNC) are shown in **Fig. 4a**. Some uTARs were co-expressed with many genes involved in cancer associated pathways, especially the P53 pathway and G2M checkpoint (**Fig. 4a**), while in kidney cancer high co-expression was observed with genes involved in P53 pathway and Epithelial-mesenchymal transition (EMT) (**Supplementary Fig. 17**). Examples of spatially co-expressed uTARs and coding genes in the H&N sample are shown in **Fig. 4b**. For example, cuTAR213507 (chr4:182814299-182820899 (+)) was co-expressed with the mRNA of *FGFR3*, a gene upregulated in H&N cancers, also on chromosome 4 (chr4:1793293-1808867-+). 24 other cancer-relevant genes located on the same chromosome with this top spatially autocorrelated cuTAR, suggesting strong evidence for a cancer-associated cuTAR. Moreover, this cuTAR contains a GWAS SNP rs1516535, an intron variant mapping to *TENM3. TENM3* is upregulated in several cancers compared to normal tissues, particularly in H&N SCCs, pancreatic adenocarcinoma, thymoma, and neuroblastoma^39^. It also serves as an integration site for the Human Papilloma Virus (HPV), causing cervical and a high proportion of Oropharyngeal cancers^40^. It is worth mentioning that the OPSCC samples utilized in this study wereHPV-16^+^. Fusions of *TENM3* gene have been reported to induce cell proliferation ^41^.

Using co-expression network analyses with hdWGCNA, we identified that cuTAR170206 formed part of a co-expression module with genes enriched for various cancer hallmarks. Additionally, it was found to be highly co-expressed with *NDRG1* **(Fig. 4c)**, which is known to be downregulated in metastatic OPSCC tumors ^42^. In this particular case, *NDRG1* was seen to be expressed only in the periphery and not in the core of the tumor (**Fig. 4c)**. This gene has been shown to have pleiotropic functions, acting as tumor promoters in some cancers while acting as a tumor suppressor in others^43^. This modulation in function could potentially be attributed to its association with interacting lncRNAs.

### Inferring functions by machine learning prediction of interaction with RBPs and colocalization

LncRNAs interact with RNA binding proteins (RBPs) to regulate mRNA and protein localization and functions^44^. Several machine learning models that have been trained on known lncRNA–RBP interactions can help predict new interactions. We applied HLPI-Ensemble^45^ to predict the interaction of the shortlisted uTARs with RNA binding proteins (RBPs). An interesting association for cuTAR215705 was observed in our in-house ST breast cancer tissues. The cell-types inferred from scType are shown (**Fig. 4d**). cuTAR215705 was predicted to interact with RNA-binding proteins (RBPs), particularly HuR (ELAVL1) and TARDBP. Moreover, it was found to be a component of the same co-expression module identified through hdWGCNA, which is enriched for TGF-***β***negative regulation and positive regulation of protein localization (**Supplementary Fig. 18a**), similar to TARDBP. Additionally, it shares this module with the gene BIRC5, which has recently been reported to interact with ELAVL1. Previous reports indicate that recombinant ELAVL1 is linked to the upregulation of BIRC5 expression, while its silencing correlates with the downregulation of both BIRC5 mRNA and protein, accompanied by increased apoptosis. Survival analyses demonstrated that increased *TTP* (ZFP36) and low BIRC5 expression predicted an overall better prognosis compared to dysregulated *TTP* and high *BIRC5*^46^. Similar trends are observed in the breast cancer data. cuTAR215705 co-localizes with *BIRC5* and *ELAVL1* in the tumor cells (**Fig. 4e**). However, little co-localization was observed with *TTP*. The co-localization was significant in the tumor region of the tissue and the absence of both the cuTAR and the two aforementioned genes (*BIRC5* and *ELAVL1*) was observed in Memory CD8^+^ T cells **(Supplementary Fig. 18b)**.

### Identification of cell-type specific lncRNAs (cuTARs) associated with response to cancer therapy

Thousands of uTARs identified in ST were consistently detected using melanoma scRNA-seq data as highlighted in **Supplementary Fig. 5**. This consistency suggests the reproducibility and validity of the identified uTARs. While Visium ST provides spatially resolved expression data, it lacks the cellular resolution provided by scRNA-seq. At single cell resolution, scRNA-seq enables the identification of individual cell-types and states, allowing for a more detailed characterization of cellular heterogeneity than in the Visium data. Integrating scRNA-seq data with ST allows assignment of spatially resolved lncRNA expression patterns to specific cell-types and subpopulations identified through scRNA-seq.

### Case study 1: Cell-type specific lncRNAs in Acral Melanoma and their associations with response to anti-PD1 immunotherapy

LncRNAs are generally specific to different cell-types and could play a role in treatment responses. To demonstrate this, a public scRNA-seq Acral Melanoma dataset was analyzed to identify cell-type specific lncRNAs and to analyze their response to anti-PD1 immunotherapy **(Supplementary Fig. 19)**. The samples were integrated, and the clusters were given cell-type identities based on marker gene expressions **(Supplementary Figures 20, 21)**. The lncRNAs differentially expressed in each cell-type were identified with edgeR **(Fig. 5a)** and expression of a subset were overlaid on the UMAP **(Fig. 5b)**. The uTARs upregulated in the tumor cells show co-expression with respective proximal protein-coding genes associated with melanoma **(Fig. 5c**). For example, we observed the co-upregulation of uTARs with pigmentation genes *PMEL* (cuTAR67350) and *MLANA* (cuTAR293520)^47^. Additionally, cuTAR280980 exhibited co-expression with ESRP1, encoding a master splicing regulator in EMT^48^, and an annotated lncRNA, *LINC00589* (also known as *TSLNC8*), implicated in diverse roles across different cancer types. In hepatocellular carcinoma, non-small cell lung cancer, and glioma, *LINC00589* inhibits proliferation, invasion, and metastasis^49^. While, in pancreatic cancer, *LINC00589* serves as an oncogene by stabilizing *CTNNB113* and is clinically valuable as an independent prognostic factor for discriminating trastuzumab responders^49^. *LINC00589* is also co-expressed with *ANXA1*, known to be upregulated in invasive melanomas ^50^. This suggests that these uTARs could be part of regulatory modules influencing tumor progression.

Up to 48.5% of lncRNAs from our in-house ST melanoma samples overlapped with the scRNA-seq melanoma data from five samples **(Fig. 5d)**. The expression of some of these overlapping uTARs across the datasets that show cell-type specific expression in the scRNA-seq melanoma samples were overlaid on the in-house ST melanoma tissues **(Figures 5e, f)**.

Random sampling and pseudo-bulking were performed on the scRNA-seq expression data pre and post anti-PD1 immunotherapy for the one sample that was available to identify lncRNAs potentially changing upon treatment. As a positive control, marker genes for melanoma were also visualized. Melanoma markers were upregulated in both cycling and non-cycling melanocytes (**Supplementary Fig. 22a)** and the proliferation markers were upregulated only in the non-cycling melanocytes (**Supplementary Fig. 22b)**. A number of uTARs were upregulated pre-treatment as compared to the same sample post treatment. For example, two lncRNAs (cuTAR288960 (chr9:22406449-22434299(+)) and cuTAR288950 (chr9:22363449-22389099(+))) were downregulated post-treatment in the melanocytes **(Figures 6a, b)**. These overlap with lncRNA annotations from other databases HSALNG0070406 (LncBook) and HSALNG0022615 (LncBook) respectively. Both of these are e-lncRNAs, overlapping with enhancers active in melanoma. 79 SNPs from GWASs associated with risk of keratinocyte cancer, BCC and other non-melanoma skin cancer are colocalized with these transcripts. On the other hand, two novel unreported potential lncRNAs cuTAR275551 (chr8:29578049-29579549(-)) and cuTAR94762 (chr15:80009299-80012299 (+)) were upregulated post-treatment in tumor cells (Figures 6a, b). The latter was co-localized with an eQTL SNP rs35541517 that affects the expression of *RASGRF1* in skin. *RASGRF1* gene fusions have been previously reported in melanoma and other cancers to induce cellular transformation and promote tumorigenesis, through activating the RAS signaling pathway^51^. LncRNA expression can be altered upon treatment and can therefore be used as markers of response. These changes in expression may also indicate that they may be playing a role in regulating oncogenesis, tumor suppression or immune resistance, given that the patient from whom this sample was collected was a poor responder and could be used as markers to indicate the same.

### Case study 2: Response to Palbociclib in Medulloblastoma PDOX models

Patient-derived orthotopic xenograft (PDOX) mouse models of human tumors are useful preclinical models to study tumors or drug effects. Treated and untreated samples from human medulloblastoma PDOX mouse models were generated as described previously^52^. These samples were analyzed for lncRNAs and those significantly differentially expressed between the treated and untreated groups were identified. Some uTARs that were expressed in the untreated human tumor region were downregulated or unexpressed in the Palbociclib treated human tumor regions **(Figures 6c, d)**. A similar trend was also observed with ONT experiments **(Supplementary Fig. 23b)** and with protein-coding oncogenes like *FOXM1, PLK1, E2F1* and *GLI2*^53^ which are oncogenes and/or involved in pathways modulated by the drug **(Supplementary Fig. 23c)**.

### uTARs as prognostic markers

LncRNAs could also be used as predictive biomarkers for the prognosis of patients with cancer^54,55^. Survival analysis was performed by dividing the samples from TCGA (360 melanoma samples) as low and high cuTAR expressing groups to see if any cuTAR could predict survival and thus be used as a prognostic marker. From the TCGA metadata, ‘Days_to_last_follow_up’ (for alive patients) and ‘days_to_death’ (for the deceased) were used to calculate survival probability. Two of the cancer-cell specific uTARs across the melanoma samples, cuTAR67350 and cuTAR293520, showed a significant difference in survival probabilities across the two groups. The low expressing group had a higher survival probability (**Supplementary Fig. 24)**. This suggests that certain spatially resolved cell-type specific lncRNAs might hold predictive value, especially at early stages. For instance, the top informative uTARs in melanoma were found to be of low or no expression in H&N cancer samples from TCGA (**Supplementary Fig. 25)** and *vice versa*. Additionally, within the relevant cancer type, patients were able to be stratified into high and low expressing groups based on the expression of these lncRNAs. Such specificity may not be achieved with lncRNAs identified from bulk data alone.

While detailed results for two cancer types were discussed, similar preliminary results for glioblastoma, kidney and breast cancers were found **(Supplementary Fig. 26)**. The cuTAR87324, upregulated in the tumor cells of kidney cancer was detected across OPSCC (validated by ONT) and skin cancers although at very low levels **(Supplementary Fig. 27)**. This could be indicative of its function specific to kidney cancers.

### A Spatial and Single Cell Pan-cancer Atlas of lncRNAs

The findings of this study have been made available through the publicly available website ‘[SPanC-Lnc](https://spanclnc.click/pages/about)’ on AWS cloud **(Fig. 7)**. The annotations and expression data can be accessed and visualized interactively. To date, this is the largest resource of pan-cancer annotations of lncRNAs with single cell and spatial context. Resources like FANTOM-CAT, lncExpDB and NONCODE provide lncRNA annotations mostly from bulk RNA-seq data of cell lines and tissues **(Table 1)**. A few recent studies have explored the utility of lncRNAs to differentiate tissue specific cell-type populations in breast cancers using single cell RNA-seq technology and highlighted the importance of the need to annotate lncRNAs with more resolution than bulk RNA-seq^16,17^. Recent studies utilizing ST data have explored the expression of annotated lncRNAs ^55,56^. These studies have incorporated methods designed to capture non-polyadenylated lncRNAs, which are often undetected or poorly detected with Visium. However, this information on a pan-cancer scale for potential novel lncRNAs is not available. This study not only provides coordinates of the identified transcripts, but also provides a comparison of expression across different cancer types and cell-types and analyzes their putative functions.

**Fig. 7:**
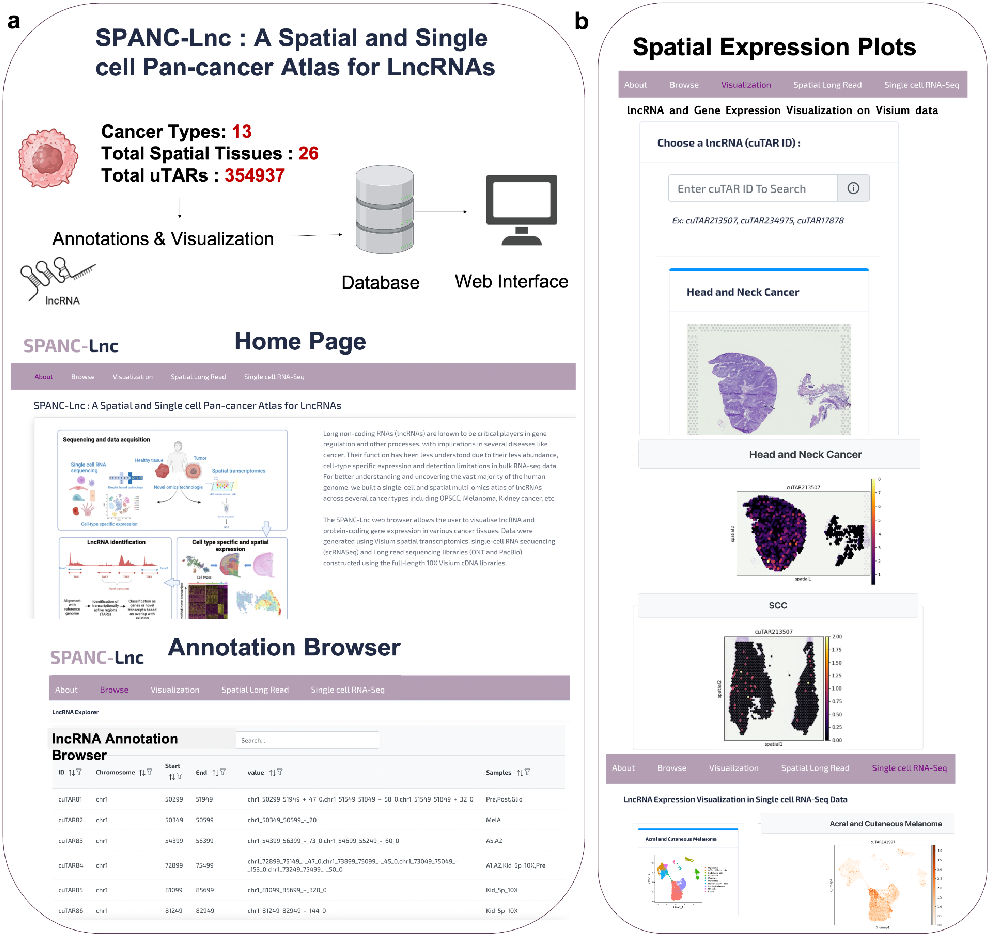
SPanC-Lnc : A Spatial and Single cell Pan-cancer Atlas for LncRNAs. **a**, All the findings of this study are available in the form of a website. The annotation browser enables perusal through lncRNA coordinates, cancer types identified, cell-type specificity, overlapping regulatory elements, etc. **b**, Visualization of the lncRNA expressions on the spatial tissues across multiple cancer types from Visium and Long Read data.

## DISCUSSION

The identification and analysis of potential novel lncRNAs from spatial and single cell datasets has provided us with valuable insights into the vast and complex landscape of gene regulation in cancers.

Our study utilized these recent technologies to profile the transcriptomes of cancer tissues, uncovering a diverse repertoire of previously unannotated lncRNAs. Through integrative analysis across multiple cancer types, we identified a subset of lncRNAs exhibiting dynamic expression patterns and tumor-specific spatial distributions, suggesting their potential roles as major regulators in tumorigenesis and cancer progression.

At a structural level, lncRNAs could be derived from enhancers, promoters, pseudogenes or transposons. LncRNAs derived from the transcription of active enhancers (e-lncRNAs) are known to participate in many cancer-associated biological processes like angiogenesis, proliferation, invasion, and metastasis 59. For example, e-lncRNA *HCCL5* when activated by ZEB1 promotes the malignant progression of hepatocellular carcinoma^60^. In colorectal cancer, e-lncRNA AC005592.2 promotes tumor progression by regulating *OLFM4* ^61^, and *LINC01488* has been shown to mediate breast cancer risk by regulating *CCND1* through estrogen signalling pathways only in tumor cells ^62^

At the genetic level, disease-associated GWAS SNPs or eQTLs could regulate nearby protein coding genes through altering the levels of co-localizing lncRNAs^63^. Further studies would be required to test these interactions as in this study we have pointed out preliminary level analysis and results. Furthermore, spatial autocorrelation with coding genes involved in several cancer-related pathways revealed coexpression of these novel lncRNAs in pathways associated with cell cycle, cellular proliferation, immune response, angiogenesis, hypoxia, and metastasis, highlighting their putative involvement in critical biological processes underlying cancer development.

Additionally, LncRNAs could also be useful in evo-devo comparative genomics. For example, identifying orthologous lncRNAs across different model organisms and studying them can help understand the functions of the human counterparts in embryonic development or other processes as highlighted in a recent study with gecko embryogenesis lncRNAs of the brain, showing they are most likely functional equivalents to human lncRNAs^64^.

While poly-adenylated lncRNAs, the main focus of this study, constitute the majority of ncRNAs, other classes of ncRNAs like circular RNAs (circRNAs)^65,66^, pre-miRNAs/miRNAs^67^, piRNAs^68,69^, and non-polyadenylated lncRNAs^70^ are also aberrantly expressed in cancers and gene regulation. Importantly, the identification of these lncRNAs in addition to the novel lncRNA panel put forth in this study would serve as a valuable resource for future investigations, providing a foundation for understanding their mechanistic roles and therapeutic potential in cancer. Single-cell sequencing data helped identify cell-type specific lncRNAs and their response to treatment in the tumor cells. Similar trends of the same lncRNAs identified across more different cancer types, could help confirm the cell-type specificity and could help serve as cell-type markers. Moreover, tumors and tissues often exhibit cellular heterogeneity. lncRNAs identified from scRNA-seq and ST data can be associated with specific subpopulations of cells, including rare cell-types that might be crucial for disease progression or prognosis. Although these may not replace the established protein-coding markers, they could complement them. Deeper analysis could be done by analyzing the survival probabilities coupled with coding genes. Additionally, our findings emphasize the significance of leveraging cutting-edge technologies to unravel the complexity of non-coding RNA landscapes, paving the way for the development of novel diagnostic markers and targeted therapies for diverse cancer types.

## ONLINE METHODS

### Datasets

#### Spatial transcriptomics datasets across different cancers

In-house and previously published datasets from fresh frozen (FF) tissues of different cancer types (Kidney^71^, Skin, SCC, BCC, Head and neck OPSCC, Medulloblastoma^52^, colorectal^72^ and Breast cancers^73^), clinical FFPE biopsies of melanoma^74^ were used in this study. Datasets from the 10X Visium FFPE probe hybridization based capture were used as the control since a predefined set of probes, targeting specific genes is used to capture the transcripts and novel ncRNAs would ideally not be captured, although off-target activity of the probes to lncRNAs has been reported^75^. Previously published in-house melanoma datasets^74^ and other publicly available datasets of ovarian and prostate cancers from the 10X Visium^75^ were used as a control as described in a previous study^75^. The coverage of the samples was an average of 100M reads captured from about 2000 spots while for the in-house kidney and breast cancer datasets it was ranging between 100-300M reads. Shown in detail are the results for the in-house spatial head and neck cancer and public melanoma scRNA-seq datasets. Similar analysis was done for all different cancer types.

#### Single-cell sequencing data

Datasets from 10X Genomics including glioblastoma, breast cancer, ovarian cancer (3’-scRNA-seq), pancreatic and kidney cancer and NSCLC (5’ scRNA-seq) were used ^76–81^.

Other previously published datasets were also used to analyze lncRNA expressions at a cell-type specific level. These included 3’-sequencing datasets of Acral and Cutaneous Melanoma from the study (PRJNA862451)^82^, PBMCs from Merkel cell carcinoma (MCC) (SRR7722937) ^83^, head and neck cancer patients (SRR13418965, SRR13419137, SRR13419168)^84^ and 5’-sequencing datasets of tissue biopsies from patients with basal cell carcinoma (BCC). This included data before and after anti-PD-1 therapy from three patients (SRX5128480, SRX5128482, SRX5128486, SRX5128489, SRX5128506, SRX5128507) out of the 12 patients in the original study (GSE123814) ^85^.

#### Detection of potential novel lncRNAs using a HMM based approach

The method described by ^21^ was adopted to identify transcriptionally active regions. The pipeline uses an R package GroHMM^86^ that utilizes a two-state hidden Markov model to classify regions in an aligned genome as transcriptionally active or not, based on the read coverage in each bin. The position sorted BAM files generated by the 10X spaceranger pipeline (spaceranger 1.3.0 and 2.0.1 using default parameters) were used as inputs to the pipeline. By default, it splits the genome into non-overlapping bins of 50bp and is called transcriptionally active if atleast three reads are detected in that bin and are labeled as TARs (Transcriptionally Active Regions). TARs found within 500 bp apart are merged into one unit. One of the limitations of this approach is that we might wrongly identify two different but adjacent transcripts as a single transcript. The regions identified are then overlapped with reference gene annotations in a strand-specific manner (reference annotations from Gencode v43 were used). The TARs overlapping with existing gene annotations (even a few base pairs of overlap is considered ‘annotated’ to account for extended gene boundaries) are labeled aTARs (annotated TARs) and the ones falling outside gene boundaries are called uTARs (unannotated TARs). We rule out the unannotated transcripts found on the opposite strand to that of an annotated transcript for this initial phase of the study. A count matrix of the TARs is generated with Drop-Seq tools DigitalExpression function^87^.

#### Analysis of sequences for coding potential, conservation and stability

To make sure that the identified transcripts are non-coding, the coding potential was analyzed using CPAT^88^. The FASTA sequences of the uTARs were extracted using bedtools getfasta. CPAT was then run using the inbuilt model for the human genome. Human coding probability (HCP) cutoff of 0.364 was used as described in the tool’s documentation. HCP >=0.364 indicates coding sequence while HCP < 0.364 indicates non-coding sequence^88^.

Further, the conservation of the sequences were calculated using the phastCons BigWig files comprising the phastCons scores for multiple alignments of 29 primate/mammalian genome sequences to the human genome GRCh38/hg38 build (Downloaded from http://hgdownload.cse.ucsc.edu/goldenPath/hg38/phastCons30way/hg38.phastCons30way.bw)^89^. The bedtools bigWigAverageOverBed function was used extract the conservation score for each specified coordinate in a bed file.

Minimum free energy (MFE) is another measure for potential functionality. In general, the lower the MFE, the more stable the secondary structure of the transcript is and hence more likely to be functional. MFE was calculated for each uTAR with RNAfold 2.6.4^90^ using the default parameters. The most stable secondary structure is predicted.

#### Classification and potential functional implications

The identified uTAR reads were overlapped with trait-associated SNPs from GWASdb^91^ and with GTEx CAVIAR eQTLs from the UCSC table browser (downloaded on 17/06/2023)^35^ with a window size of 10kb using bedtools to propose potential genetic level regulation based on co-localizing features.

The identified uTARs were overlapped with enhancers from the EnhancerAtlas^33^ and TSS data from FANTOM for window sizes ranging from 1kb-1Mb. These can also provide important clues about the regulatory mechanisms and potential functions of these transcripts, helping to shed light on the complex regulatory networks that govern gene expression.

#### Identification of cancer-specific lncRNAs

The tumor regions in each tissue were identified using annotations by a pathologist and using gene expression profiles. The loupe browser from 10X Genomics was used to label the barcodes as cancerous and normal and the annotations were exported and used for further analysis. The uTARs differentially expressed across these two annotated clusters were identified using Seurat v4.3 processed using SCTransform (for FindAllMarkers function, the parameters min.pct was set to 0.1 and logfc.threshold varied between 0.1-0.25 depending on the sample analyzed. The percentage of spots in the cancerous *versus* normal regions expressing the selected uTARs were also calculated.

Few uTARs from two samples of the Head and Neck cancer dataset were shortlisted to be the uTARs of interest that could be the top potential functional candidates. They were chosen such that (i) they are detected in all samples (ii) they are longer than 1000 bp (iii) they are novel or represented in public lncRNA databases (iv) they are differentially expressed in the cancerous region of the tissue. To be more confident about the chosen regions, we also checked their coverages in bulk RNA-seq datasets from TCGA and the scRNA-seq dataset of HNC and checked if they were detected. The bigwig RNA-seq coverage files for 96 random TCGA samples were downloaded using recount. The uTAR coverages were extracted from these files using bigWigAverageOverBed for the specified boundaries in a .bed file. TMM normalization was performed on the raw coverage (counts) using edgeR. By analyzing coverage, gene expression levels can be inferred. Regions with higher coverage indicate higher expression levels of the corresponding uTARs.

#### Confirmation of detected signals using Long-read sequencing

##### Oxford Nanopore

Long-read sequencing with Oxford Nanopore technology was performed using the 10X Visium biotinylated 3’ cDNA libraries to validate the expression of the identified potential lncRNAs using Promethion flow cells. SQK-LSK110 and EXP-NBD kits from ONT was used to generate the libraries for the Head and Neck sample C (HNC). Sequence runs were generated using super-accurate basecalling setting with MinKNOW version 22.12.5 and Guppy version 6.4.6 (flow cell type: FLO-PRO002). The nanopore run generated ∼16 million reads. To demultiplex the reads based on the spatial barcode, scNanoGPS was used. Some changes were made to the default values of the pipeline with respect to the scan region, which was set to 1500bp since the 3’ adaptors were found as many bases away due to longer chimeras. The long non-coding RNAs were then identified using a modified version of the uTAR pipeline, since scNanoGPS generates individual alignment files for each spatial barcode identified, rather than an individual BAM file with reads tagged with spatial/cell and UMI barcodes. Featurecounts was used to generate the uTAR expression matrix. GTF annotations of the uTARs from the corresponding 10X Visium dataset was also used to generate the count matrix. The identified spatial barcodes were matched with those from the 10X Visium barcode list from the tissue_positions_list.csv in the spaceranger output of the corresponding Visium data and only those with tissues placed on them were retained. The coordinates of the matched barcodes were also extracted from the aforementioned CSV file. The expression of some potential candidates were visualized using stlearn^92^ and was compared to that of the previously generated short-read data.

Further, the experiment was also performed for skin cancer samples. SQK-NBD-114.24 kit (Native barcoding kit 14) to prepare libraries for the SCC and BCC samples, which were sequenced on PromethION P24 device using R10 flow cell (FLO-PRO114M). Sequence bases (raw data) were called using PromethION software release 23.11.4 (minknow-core-promethion 5.8.3, dorado version 7.2.13). Re-basecalling was done using dorado version 0.5.1. Similar downstream analysis using scNanoGPS and uTAR visualization were performed.

##### PacBio

The full length Visium cDNA libraries were used for HiFi Sequencing. The procedure for Preparing MAS-Seq libraries using MAS-Seq for 10x Single Cell 3’ kit (PN:102-678-600-REV03) is as follows. After the steps of cDNA amplification, cleanup and QC of 10X Visium, 225 pM cDNA per library with concentration of 23 ng/ul were pooled for sequencing. Firstly, the TSO priming artifacts during cDNA synthesis were removed using biotinylated primers. Next, DNA fragments containing orientation-specific MAS segmentation adapter sequences were generated by performing 16 parallel cDNA amplification reactions. MAS enzyme was used to create single-stranded extensions to enable directional assembly of cDNA segments into a linear array. After DNA damage repair and nuclease treatment, the cDNA clean-up was done with 1.2X SMRTbell beads.

Further, the pbcromwell workflow of SMRTlink Tools was used for to process the HiFi BAM files. With the aligned genome file from Minimap2, the uTAR pipeline was run and the uTAR expression was overlaid on the tissues and compared with that of the Visium.

##### Quantitative Reverse Transcription Polymerase Chain Reaction

Six cDNA samples from the 10X Visium libraries including the Head and Neck Samples (HNB, HNC), SCC, BCC and the Colorectal cancer samples (Colorectal Primary Tumor - CP and metastasized tumor - CM) were tested for seven potential lncRNAs **(Table)**. *GAPDH*, a housekeeping gene and *KRT18*, an epithelial marker were used as positive controls. Two negative controls NEG1 and NEG2 were designed to target genomic regions around the centromere that do not code for RNA, hence controls for any genomic DNA contamination. H2O was used as a non-template negative control. 12 primers were designed to target 7 uTARs. The primers were designed in such a way that they did not have any off-targets using the sequence information of the HNC sample. qPCR Master Mix was prepared for each sample following the KAPA SYBR FAST qPCR Kit guide to allow each well to contain 1μL of 10ng/μL cDNA, 5μL KAPA SYBR® FAST qPCR Master Mix (2X) (Roche), 0.2μL ROX™ Low Reference Dye (50X) (Roche) and 3.4μL of RNAse-free water. As GAPDH is an endogenous control, the template for this primer was further diluted 1:8 to ensure it is within a comparable quantifiable range.

9.6μL of the qPCR Master Mix was loaded into each well on the 96-well qRT-PCR plate before 0.4μL of 10nM the respective lncRNA primer pairs were added. The qPCR reaction was performed with the following protocol using the ViiA7 96-well Real-Time PCR System with High Resolution Melt (Applied Biosystems); initial denaturation (98°C for 3 minutes), 2-step amplification (98°C for 5 seconds and 63°C for 30 seconds) for a total of 35 cycles and the melting curve method.

The expression was calculated using the formula C_KRT18_.CT_KRT18_=C_cuTAR_.CT_cuTAR_. The CT was normalized with that of one of the positive controls *KRT18* such that C_KRT18_ = 1. The cuTAR expression was calculated as C_cuTAR_ = 1.CT_KRT18_/CT_cuTAR_. The average was used for two primers targeting a same cuTAR. The expression was visualized on a heatmap **(Supplementary Fig. 14)**.

##### Spatial autocorrelation of lncRNAs with genes relevant to cancer

SpatialDE^93^ was used to identify spatially variable features. The significant spatially variable uTARs and the top chosen candidates chosen for the Head and neck cancer samples were further used to measure pairwise spatial autocorrelation with coding genes using Moran’s Index using python scripts. Several hallmark genes relevant to cancer were retrieved from GSEA-MsigDB^94^ and were used for the analysis. The spots showing (i) high expression of both the gene and the uTAR were categorized as HH, (ii) spots with gene expression and no uTAR expression as LH, (iii) ones with uTAR expression and lack of gene expression as HL (iv) no expression of both features as LL and (v) ns for spots with insignificant Moran’s I. The uTARs showing high spatial correlation (more than 50 HH spots) with genes of different hallmarks were calculated to identify the more functionally relevant uTARs and the pathways involved. The Moran’s I was calculated using the formula below

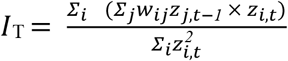

Here, *I*_T_ is the calculated Bivariate Moran’s Index Bivariate Spatial Correlation for expression values at two points: lncRNA (t) and gene (t-1) across all spots with central spots *i*, and their neighboring spots *j*. Spatial weights (w_*ij*_) represent the strength of spatial interaction or proximity between spatial units *i* and *j*. These weights are defined based on contiguity (e.g., sharing a border). With this, *I*_*T*_ is 1 if all neighboring spots z_j,t-1_ have the same value of that gene at *z*_*i,t*_. Therefore, *I*_*T*_ = 1 indicates that the predicted value of a gene with a high spatial autocorrelation is accurate (based on the values of neighboring spots). In another word, *I*_*T*_ measures the degree of *spatial* correlation between observed values in neighboring spots and the predicted values of central spots.

##### Interaction with RBPs and colocalization

Machine learning models that have been trained using known lncRNA-protein interactions can be used to predict interaction of novel lncRNAs with proteins, adding another layer or potential functionality. HLPI-Ensemble which adopts the ensemble strategy based on three mainstream machine learning algorithms of Support Vector Machines (SVM), Random Forests (RF) and Extreme Gradient Boosting (XGB) was used to predict the interaction of top uTARs with RNA binding proteins (RBPs). Further the spatial colocalization of the uTARs with their interacting RBP partners and with the genes with reported associated with those RBPs was visualized.

##### Identification of cell-type specific lncRNAs and their response to therapy

The melanoma scRNA-seq dataset from the study (PRJNA862451)^82^ was used to identify cell-type specific lncRNAs and their response to therapy. The dataset included acral and cutaneous melanoma samples. The coding gene count matrices 4 acral and 1 cutaneous melanoma samples (10X Chromium V2 chemistry) were integrated using the Seurat workflow and clustering was performed. Cell-type markers as described in the original study were used to annotate the clusters. Melanocytes (*MITF, PMEL, TYR, DCT, MLANA, PMEL, APOC1, S100A1*) Cycling melanoma cells (*UBE2C, NUSAP1, MKI67, CENPF*), Endothelial cells (*VWF, PECAM1*), Fibroblasts (*COL3A1, COL1A2, COL1A1, LUM*), CD8+ T-cells (*CD8A, HAVCR2, LAG3, PD1, TIGIT, CTLA4, HOPX*), CD4+ T-cells (*CD4, FOXP3, IL2*), Monocytes (*CD14, LYZ, CD74, CD68, CD79A*), B cells (*MS4A1*), NK cells (*GNLY, FGFBP2, FCGR3A, KLRD1, KLRF1*), NK-like T-cells (*CD3E, CD3D, GZMB, XCL2, IFNG, CCL4, NKG7, GZMA, GZMK*) and macrophages (*CD68, CD163, CD14, CD11b, CD206, CD80, CD86, CD16, CD64, CCL18, CD115, CD11c, CD32, HLA-DR, MRC1, MSR1, GCA, Pf4*) were identified.

The cell barcode and its corresponding cell-type identity were used as metadata for the differential expression analysis of the potential lncRNAs. DE analysis was performed by the pseudo-bulk approach using edgeR. The model was built as design<-model.matrix(∼celltype+sample) to identify cell-type specific lncRNAs accounting for sample batch effects.

Further there was only one acral melanoma sample for which data was collected before and after anti-PD1 treatment. The differentially expressed genes and lncRNAs in the tumor cells with respect to the other cells were identified. 10 pseudo-replicates of 60 cells per-cell-type were created and their raw expression across cell-types was visualized. As a positive control, marker genes for melanoma were also visualized.

##### Response to drug in Medulloblastoma PDOX models

In-house Visium Medulloblastoma data as described by^52^ were also analyzed. Briefly, Medulloblastoma (SHH MB) tissue from a 4.9-year-old patient was used to generate PDOX line by implanting the tumor cells in the cerebellum of immunocompromised NSG mice within hours of surgical removal from the patient and propagating them from mouse to mouse exclusively without in vitro passaging as previously described. Two of the tumor bearing mice were treated with Palbociclib hydrochloride (Pfizer) and the rest were untreated. These treated and untreated samples were analyzed for lncRNAs in this study and those significantly differentially expressed between the treated and untreated groups were identified. The raw Visium data were processed using a human-mouse hybrid reference since the tissue comprised of the human tumor and the mouse brain tissues.

##### Interactive Website for annotations and visualization

The findings of this study have been made available for users to browse through the annotations and to visualize coding gene and cuTAR expression on tissue images. The Angular 13.3 web site is hosted using AWS Amplify. It uses a Github repository for version control, AWS Code Build to generate the production artefacts (HTML, CSS, and JavaScript), and uses AWS Cloudfront CDN (https://aws.amazon.com/cloudfront/features/) to distribute these globally. Libraries used include: TypeScript 4.6 and Node 10.8.1. The backend web service is running the Django 5.0 web framework, on Python 3.10, using the slim version of Debian Bookworm (https://hub.docker.com/_/debian). This is deployed on AWS Lambda instances provisioned with 4.5 GB of RAM connected to an AWS EFS (Elastic File System) file system for accessing HDF5 and SQLite 3 files used during querying. Python libraries used include: AnnData 0.10, h5py 3.11.0, numba 0.59.1, numpy 1.26.4, pandas 2.2.2, scanpy 1.10.1, scipy 1.13.1, and seaborn 0.13.2. For visualization of expressions, python scripts were used in the backend with preloaded anndata objects for each sample (.h5ad files).

## DATA AVAILABILITY

The annotations generated in this study is available on the website SPanC-Lnc. All the scripts used for the analysis and visualizations can be accessed on Github at https://github.com/GenomicsMachineLearning/SPanc_Lnc_PanCancer_LncRNA_Atlas.git. The count matrices will be published to UQ eSPACE. The raw data of the unpublished in-house data will be submitted to the European Genome-Phenome Archive and will be accessible upon request.

## SUPPLEMENTARY DATA

Supplementary Data are available online.

## AUTHOR CONTRIBUTIONS

P. Prakrithi: Formal analysis, Methodology, Visualization, Validation, Writing—original draft. Tuan Vo: Experiments. Hani Vu: qPCR experiments. Andrew Newman— Website hosting on AWS cloud, Jazmina Gonzalez Cruz: Writing—review & editing. Ishaan Gupta: Conceptualization, Writing—review & editing. Quan Nguyen: Conceptualization, Writing—review & editing.

## ACKNOWLEDGEMENT

We acknowledge Subash Rai and Shivangi Wani for their contribution to the Nanopore and PacBio experiments respectively. We also thank B. Sathish Kumar for building the SPanC-Lnc website and Xingliang from PacBio team for helping setup SMRTlink. We would like to thank Laura Grice and Juliet French for their critical review of the manuscript and their valuable suggestions.

## ETHICS APPROVAL STATEMENT

All samples were approved for research under ethical approval numbers 2018000165 and 2017000318 by the University of Queensland’s Human Research Ethics Committees.

## Additional information

Supplementary information The online version contains supplementary material available at Supplementary_final.pdf. Correspondence and requests for materials should be addressed to Quan Nguyen.

## REFERENCES

1. Boland, C. R. Non-coding RNA: It’s Not Junk. Dig. Dis. Sci. 62, 1107–1109 (2017).

2. Niderla-Bielińska, J., Jankowska-Steifer, E. & Włodarski, P. Non-Coding RNAs and Human Diseases: Current Status and Future Perspectives. Int. J. Mol. Sci. 24, 11679 (2023).

3. Fang, Y. & Fullwood, M. J. Roles, Functions, and Mechanisms of Long Non-coding RNAs in Cancer. Genomics Proteomics Bioinformatics 14, 42–54 (2016).

4. Zhang, X. et al. Mechanisms and Functions of Long Non-Coding RNAs at Multiple Regulatory Levels. Int. J. Mol. Sci. 20, 5573 (2019).

5. Statello, L., Guo, C.-J., Chen, L.-L. & Huarte, M. Gene regulation by long non-coding RNAs and its biological functions. Nat. Rev. Mol. Cell Biol. 22, 96–118 (2021).

6. Wang, L. et al. CRISPR-Cas13d screens identify KILR, a breast cancer risk-associated lncRNA that regulates DNA replication and repair. Mol. Cancer 23, 101 (2024).

7. Wang, Y. et al. LncRNA-encoded polypeptide ASRPS inhibits triple-negative breast cancer angiogenesis. J. Exp. Med. 217, jem.20190950 (2020).

8. Jin, H. et al. lncRNA and breast cancer: Progress from identifying mechanisms to challenges and opportunities of clinical treatment. Mol. Ther. Nucleic Acids 25, 613–637 (2021).

9. Qian, Y., Shi, L. & Luo, Z. Long Non-coding RNAs in Cancer: Implications for Diagnosis, Prognosis, and Therapy. Front. Med. 7, 612393 (2020).

10. Lemos, A. E. G., Matos, A. da R., Ferreira, L. B. & Gimba, E. R. P. The long non-coding RNA PCA3: an update of its functions and clinical applications as a biomarker in prostate cancer. Oncotarget 10, 6589–6603 (2019).

11. Modi, A. et al. Integrative Genomic Analyses Identify LncRNA Regulatory Networks across Pediatric Leukemias and Solid Tumors. Cancer Res. 83, 3462–3477 (2023).

12. Tan, B.-S. et al. LncRNA NORAD is repressed by the YAP pathway and suppresses lung and breast cancer metastasis by sequestering S100P. Oncogene 38, 5612–5626 (2019).

13. Puvvula, P. K. et al. Long noncoding RNA PANDA and scaffold-attachment-factor SAFA control senescence entry and exit. Nat. Commun. 5, 5323 (2014).

14. Park, E.-G., Pyo, S.-J., Cui, Y., Yoon, S.-H. & Nam, J.-W. Tumor immune microenvironment lncRNAs. Brief. Bioinform. 23, bbab504 (2021).

15. Yang, L., Duff, M. O., Graveley, B. R., Carmichael, G. G. & Chen, L.-L. Genomewide characterization of non-polyadenylated RNAs. Genome Biol. 12, R16 (2011).

16. Bitar, M. et al. Redefining normal breast cell populations using long noncoding RNAs. Nucleic Acids Res. 51, 6389–6410 (2023).

17. Pinkney, H. R., Black, M. A. & Diermeier, S. D. Single-Cell RNA-Seq Reveals Heterogeneous lncRNA Expression in Xenografted Triple-Negative Breast Cancer Cells. Biology 10, 987 (2021).

18. Spatial transcriptome analysis of long non-coding RNAs reveals tissue specificity and functional roles in cancer. J. Zhejiang Univ. Sci. B 24, 15–31 (2023).

19. Lv, D. et al. LncSpA: LncRNA Spatial Atlas of Expression across Normal and Cancer Tissues. Cancer Res. 80, 2067–2071 (2020).

20. Weirick, T. et al. The identification and characterization of novel transcripts from RNA-seq data. Brief. Bioinform. 17, 678–685 (2016).

21. Wang, M. F. Z. et al. Uncovering transcriptional dark matter via gene annotation independent single-cell RNA sequencing analysis. Nat. Commun. 12, 2158 (2021).

22. Xu, K. et al. Pan-cancer characterization of expression and clinical relevance of m6A-related tissue-elevated long non-coding RNAs. Mol. Cancer 20, 31 (2021).

23. Li, Y. et al. Pan-cancer characterization of immune-related lncRNAs identifies potential oncogenic biomarkers. Nat. Commun. 11, 1000 (2020).

24. Isaev, K. et al. Pan-cancer analysis of non-coding transcripts reveals the prognostic onco-lncRNA HOXA10-AS in gliomas. Cell Rep. 37, 109873 (2021).

25. Luo, Y., Morgan, S. L. & Wang, K. C. PICSAR: Long Noncoding RNA in Cutaneous Squamous Cell Carcinoma. J. Invest. Dermatol. 136, 1541–1542 (2016).

26. Wang, Q. et al. LncRNA TINCR impairs the efficacy of immunotherapy against breast cancer by recruiting DNMT1 and downregulating MiR-199a-5p via the STAT1– TINCR-USP20-PD-L1 axis. Cell Death Dis. 14, 76 (2023).

27. Lu, D. et al. The long noncoding RNA TINCR promotes breast cancer cell proliferation and migration by regulating OAS1. Cell Death Discov. 7, 1–16 (2021).

28. Lin, Z.-B. et al. Long Noncoding RNA KCNQ1OT1 is a Prognostic Biomarker and mediates CD8+ T cell exhaustion by regulating CD155 Expression in Colorectal Cancer. Int. J. Biol. Sci. 17, 1757–1768 (2021).

29. Mini, E. et al. RNA sequencing reveals PNN and KCNQ1OT1 as predictive biomarkers of clinical outcome in stage III colorectal cancer patients treated with adjuvant chemotherapy. Int. J. Cancer 145, 2580–2593 (2019).

30. Ma, L., Bajic, V. B. & Zhang, Z. On the classification of long non-coding RNAs. RNA Biol. 10, 924–933 (2013).

31. Li, J. & Liu, C. Coding or Noncoding, the Converging Concepts of RNAs. Front. Genet. 10, 496 (2019).

32. Pan, J. et al. Functional Micropeptides Encoded by Long Non-Coding RNAs: A Comprehensive Review. Front. Mol. Biosci. 9, 817517 (2022).

33. Gao, T. & Qian, J. EnhancerAtlas 2.0: an updated resource with enhancer annotation in 586 tissue/cell types across nine species. Nucleic Acids Res. 48, D58–D64 (2020).

34. The GTEx Consortium atlas of genetic regulatory effects across human tissues. Science 369, 1318–1330 (2020).

35. Lonsdale, J. et al. The Genotype-Tissue Expression (GTEx) project. Nat. Genet. 45, 580–585 (2013).

36. Chen, E. Y. et al. Enrichr: interactive and collaborative HTML5 gene list enrichment analysis tool. BMC Bioinformatics 14, 128 (2013).

37. The Cancer Genome Atlas - Citing TCGA - National Cancer Institute. https://www.cancer.gov/about-nci/organization/ccg/research/structural-genomics/tcga/using-tcga/citing-tcga (2019).

38. Gao, R. et al. Delineating copy number and clonal substructure in human tumors from single-cell transcriptomes. Nat. Biotechnol. 39, 599–608 (2021).

39. Peppino, G. et al. Teneurins: Role in Cancer and Potential Role as Diagnostic Biomarkers and Targets for Therapy. Int. J. Mol. Sci. 22, 2321 (2021).

40. Jang, M. K., Shen, K. & McBride, A. A. Papillomavirus genomes associate with BRD4 to replicate at fragile sites in the host genome. PLoS Pathog. 10, e1004117 (2014).

41. Hiwatari, M. et al. Abstract 6734: Identification of the novel TENM3-ALK fusion in an AYA case with ALK rearranged neuroblastoma. Cancer Res. 83, 6734 (2023).

42. de Lima, J. M. et al. NDRG1 deficiency is associated with regional metastasis in oral cancer by inducing epithelial-mesenchymal transition. Carcinogenesis 41, 769–777 (2020).

43. Joshi, V., Lakhani, S. R. & McCart Reed, A. E. NDRG1 in Cancer: A Suppressor, Promoter, or Both? Cancers 14, 5739 (2022).

44. Huang, H., Li, L. & Wen, K. Interactions between long non-coding RNAs and RNA-binding proteins in cancer. Oncol. Rep. 46, 256 (2021).

45. Hu, H. et al. HLPI-Ensemble: Prediction of human lncRNA-protein interactions based on ensemble strategy. RNA Biol. 15, 797–806 (2018).

46. Al-Yahya, S. et al. Post-transcriptional regulation of BIRC5/survivin expression and induction of apoptosis in breast cancer cells by tristetraprolin. RNA Biol. 21, 1–15 (2024).

47. Kawakami, A. & Fisher, D. E. The master role of microphthalmia-associated transcription factor in melanocyte and melanoma biology. Lab. Invest. 97, 649–656 (2017).

48. Yao, J. et al. Altered Expression and Splicing of ESRP1 in Malignant Melanoma Correlates with Epithelial–Mesenchymal Status and Tumor-Associated Immune Cytolytic Activity. Cancer Immunol. Res. 4, 552–561 (2016).

49. Bai, W. et al. LINC00589-dominated ceRNA networks regulate multiple chemoresistance and cancer stem cell-like properties in HER2+ breast cancer. Npj Breast Cancer 8, 1–19 (2022).

50. Delorme, S. et al. New insight into the role of ANXA1 in melanoma progression: involvement of stromal expression in dissemination. Am. J. Cancer Res. 11, 1600–1615 (2021).

51. Hunihan, L. et al. RASGRF1 Fusions Activate Oncogenic RAS Signaling and Confer Sensitivity to MEK Inhibition. Clin. Cancer Res. Off. J. Am. Assoc. Cancer Res. 28, 3091–3103 (2022).

52. Vo, T. et al. Spatial transcriptomic analysis of Sonic hedgehog medulloblastoma identifies that the loss of heterogeneity and promotion of differentiation underlies the response to CDK4/6 inhibition. Genome Med. 15, 29 (2023).

53. Huang, D. et al. GLI2 promotes cell proliferation and migration through transcriptional activation of ARHGEF16 in human glioma cells. J. Exp. Clin. Cancer Res. CR 37, 247 (2018).

54. Yu, P., Ye, J., Zhao, S. & Cai, Y. lncRNAs are potential prognostic markers in patients with nasopharyngeal carcinoma in China: A systematic review and meta-analysis. Mol. Clin. Oncol. 20, 1–13 (2024).

55. Arriaga-Canon, C. et al. The use of long non-coding RNAs as prognostic biomarkers and therapeutic targets in prostate cancer. Oncotarget 9, 20872–20890 (2018).

56. Hon, C.-C. et al. An atlas of human long non-coding RNAs with accurate 5′ ends. Nature 543, 199–204 (2017).

57. Li, Z. et al. LncExpDB: an expression database of human long non-coding RNAs. Nucleic Acids Res. 49, D962–D968 (2021).

58. Lianhe, Z. et al. NONCODEV6: An updated database dedicated to long non-coding RNA annotation in both animals and plants. Nucleic Acids Res. 49, (2020).

59. García-Padilla, C. et al. Molecular Mechanisms of lncRNAs in the Dependent Regulation of Cancer and Their Potential Therapeutic Use. Int. J. Mol. Sci. 23, 764 (2022).

60. Peng, L. et al. Super-Enhancer-Associated Long Noncoding RNA HCCL5 Is Activated by ZEB1 and Promotes the Malignancy of Hepatocellular Carcinoma. Cancer Res. 79, 572–584 (2019).

61. Yan, L., Chen, H., Tang, L., Jiang, P. & Yan, F. Super-enhancer-associated long noncoding RNA AC005592.2 promotes tumor progression by regulating OLFM4 in colorectal cancer. BMC Cancer 21, 187 (2021).

62. Bjørklund, S. S. et al. Subtype and cell type specific expression of lncRNAs provide insight into breast cancer. Commun. Biol. 5, 1–14 (2022).

63. Castellanos-Rubio, A. & Ghosh, S. Functional Implications of Intergenic GWAS SNPs in Immune-Related LncRNAs. in Long Noncoding RNA: Mechanistic Insights and Roles in Inflammation (ed. Carpenter, S.) 147–160 (Springer International Publishing, Cham, 2022). doi:10.1007/978-3-030-92034-0_8.

64. Olazagoitia-Garmendia, A., Senovilla-Ganzo, R., García-Moreno, F. & Castellanos-Rubio, A. Functional evolutionary convergence of long noncoding RNAs involved in embryonic development. Commun. Biol. 6, 1–11 (2023).

65. Kristensen, L. S., Jakobsen, T., Hager, H. & Kjems, J. The emerging roles of circRNAs in cancer and oncology. Nat. Rev. Clin. Oncol. 19, 188–206 (2022).

66. Yarmishyn, A. A. et al. Circular RNAs Modulate Cancer Hallmark and Molecular Pathways to Support Cancer Progression and Metastasis. Cancers 14, 862 (2022).

67. MacFarlane, L.-A. & Murphy, P. R. MicroRNA: Biogenesis, Function and Role in Cancer. Curr. Genomics 11, 537–561 (2010).

68. Zhang, Q. et al. The epigenetic regulatory mechanism of PIWI/piRNAs in human cancers. Mol. Cancer 22, 45 (2023).

69. Yao, J. et al. PIWI-interacting RNAs in cancer: Biogenesis, function, and clinical significance. Front. Oncol. 12, (2022).

70. Zhang, Y., Yang, L. & Chen, L.-L. Life without A tail: new formats of long noncoding RNAs. Int. J. Biochem. Cell Biol. 54, 338–349 (2014).

71. Raghubar, A. M. et al. High risk clear cell renal cell carcinoma microenvironments contain protumour immunophenotypes lacking specific immune checkpoints. Npj Precis. Oncol. 7, 1–9 (2023).

72. Kawamata, F. et al. Copy number profiles of paired primary and metastatic colorectal cancers. Oncotarget 9, 3394–3405 (2017).

73. Wu, S. Z. et al. A single-cell and spatially resolved atlas of human breast cancers. Nat. Genet. 53, 1334–1347 (2021).

74. Vo, T. et al. Benchmarking Robust Spatial Transcriptomics Approaches to Capture the Molecular Landscape and Pathological Architecture of Archived Cancer Tissues. (2023). doi:10.1101/2023.02.11.527941.

75. Prakrithi, P., Juwayria, Jain D., Malik, P. S. & Gupta, I. Caution towards spurious off-target signal in 10X Visium spatial transcriptomics assay from potential lncRNAs. Brief. Bioinform. bbad031 (2023) doi:10.1093/bib/bbad031.

76. Human Ovarian Tumor (FF) (v2, 150 × 150), Single Cell Immune Profiling Dataset by Cell Ranger 7.0.0, 10x Genomics, (2022, May 14).

77. Human Invasive Ductal Carcinoma (3’ v3.1, 150 × 150), Single Cell Gene Expression Dataset by Cell Ranger 6.0.0, 10x Genomics, (2021, March 31).

78. Human Glioblastoma Multiforme (3’ v3.1, 150 × 150), Single Cell Gene Expression Dataset by Cell Ranger 6.0.0, 10x Genomics, (2021, March 31).

79. Human Kidney Tumor (FF) (5’ v2, 150 × 150), Single Cell Immune Profiling Dataset by Cell Ranger 7.0.0, 10x Genomics, (2022, May 14).

80. Pancreatic Tumor (FF) (5’ v2, 150 × 150), Single Cell Immune Profiling Dataset by Cell Ranger 7.0.0, 10x Genomics, (2022, May 14).

81. NSCLC Tumor (F) (v2, 150 × 150), 5’ Single Cell Immune Profiling Dataset by Cell Ranger 2.2.0, 10x Genomics, (2018, August 1).

82. Zhang, C. et al. A single-cell analysis reveals tumor heterogeneity and immune environment of acral melanoma. Nat. Commun. 13, 7250 (2022).

83. Paulson, K. G. et al. Acquired cancer resistance to combination immunotherapy from transcriptional loss of class I HLA. Nat. Commun. 9, 3868 (2018).

84. Kürten, C. H. L. et al. Investigating immune and non-immune cell interactions in head and neck tumors by single-cell RNA sequencing. Nat. Commun. 12, 7338 (2021).

85. Yost, K. E. et al. Clonal replacement of tumorspecific T cells following PD-1 blockade. Nat. Med. 25, 1251–1259 (2019).

86. Chae, M., Danko, C. G. & Kraus, W. L. groHMM: a computational tool for identifying unannotated and cell type-specific transcription units from global run-on sequencing data. BMC Bioinformatics 16, 222 (2015).

87. Bageritz, J. & Raddi, G. Single-Cell RNA Sequencing with Drop-Seq. Methods Mol. Biol. Clifton NJ 1979, 73–85 (2019).

88. Wang, L. et al. CPAT: Coding-Potential Assessment Tool using an alignment-free logistic regression model. Nucleic Acids Res. 41, e74 (2013).

89. Pollard, K. S., Hubisz, M. J., Rosenbloom, K. R. & Siepel, A. Detection of nonneutral substitution rates on mammalian phylogenies. Genome Res. 20, 110–121 (2010).

90. Lorenz, R. et al. ViennaRNA Package 2.0. Algorithms Mol. Biol. 6, 26 (2011).

91. Li, M. J. et al. GWASdb: a database for human genetic variants identified by genome-wide association studies. Nucleic Acids Res. 40, D1047–D1054 (2012).

92. Pham, D. et al. stLearn: integrating spatial location, tissue morphology and gene expression to find cell types, cell-cell interactions and spatial trajectories within undissociated tissues. bioRxiv 2020.05.31.125658 (2020) doi:10.1101/2020.05.31.125658.

93. Svensson, V., Teichmann, S. A. & Stegle, O. SpatialDE: identification of spatially variable genes. Nat. Methods 15, 343–346 (2018).

94. Subramanian, A. et al. Gene set enrichment analysis: A knowledge-based approach for interpreting genome-wide expression profiles. Proc. Natl. Acad. Sci. 102, 15545–15550 (2005).

