## Supplementary material for "Unraveling lncRNA Diversity at a Single Cell Resolution and in a Spatial Context across Different Cancer Types": Supplementary_material-compressed.pdf

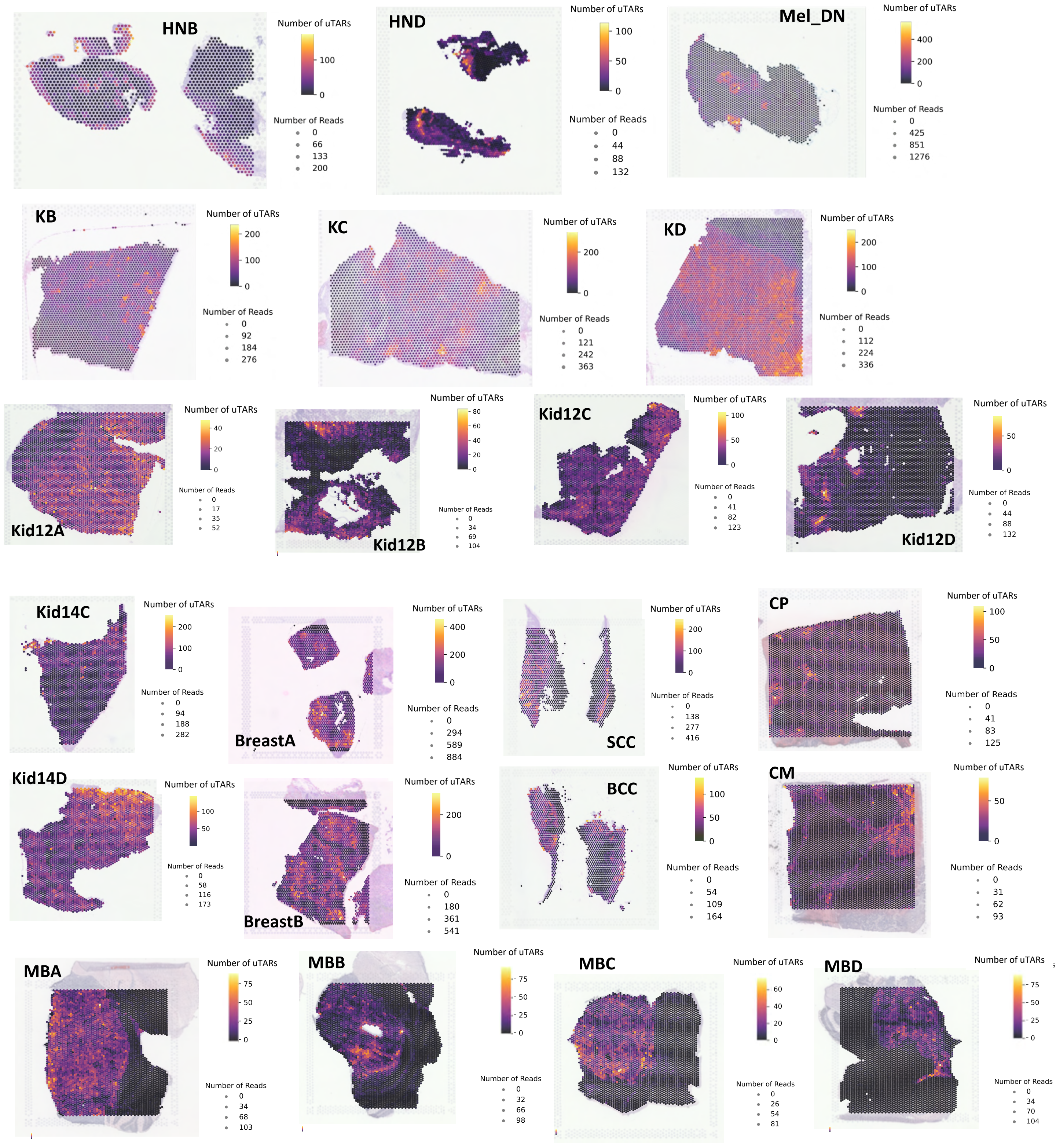


**Supplementary Fig. 1:** uTAR counts overlaid on the tissue for all in-house samples

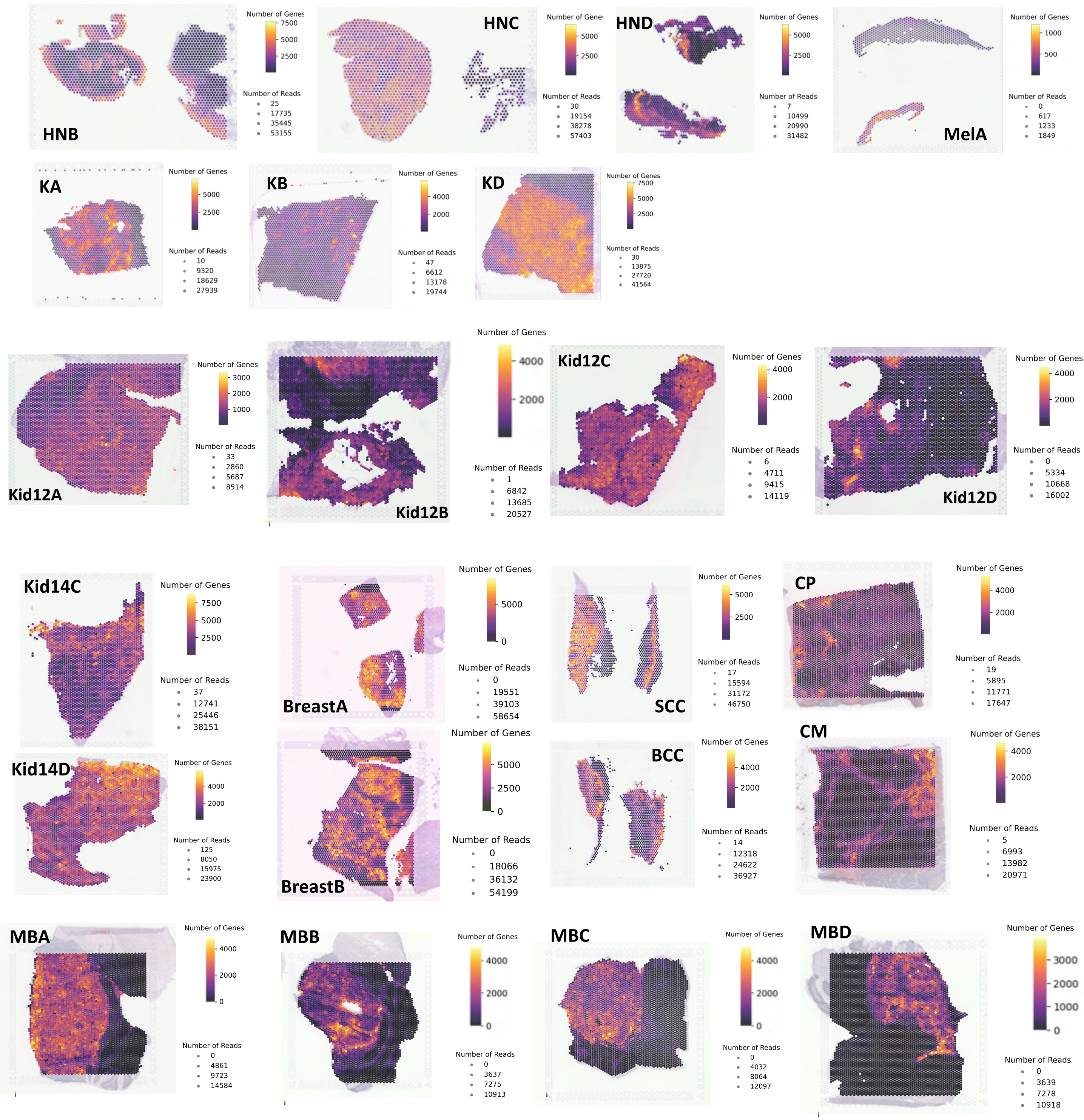


**Supplementary Fig. 2:** Coding gene counts overlaid on the tissue for all in-house samples

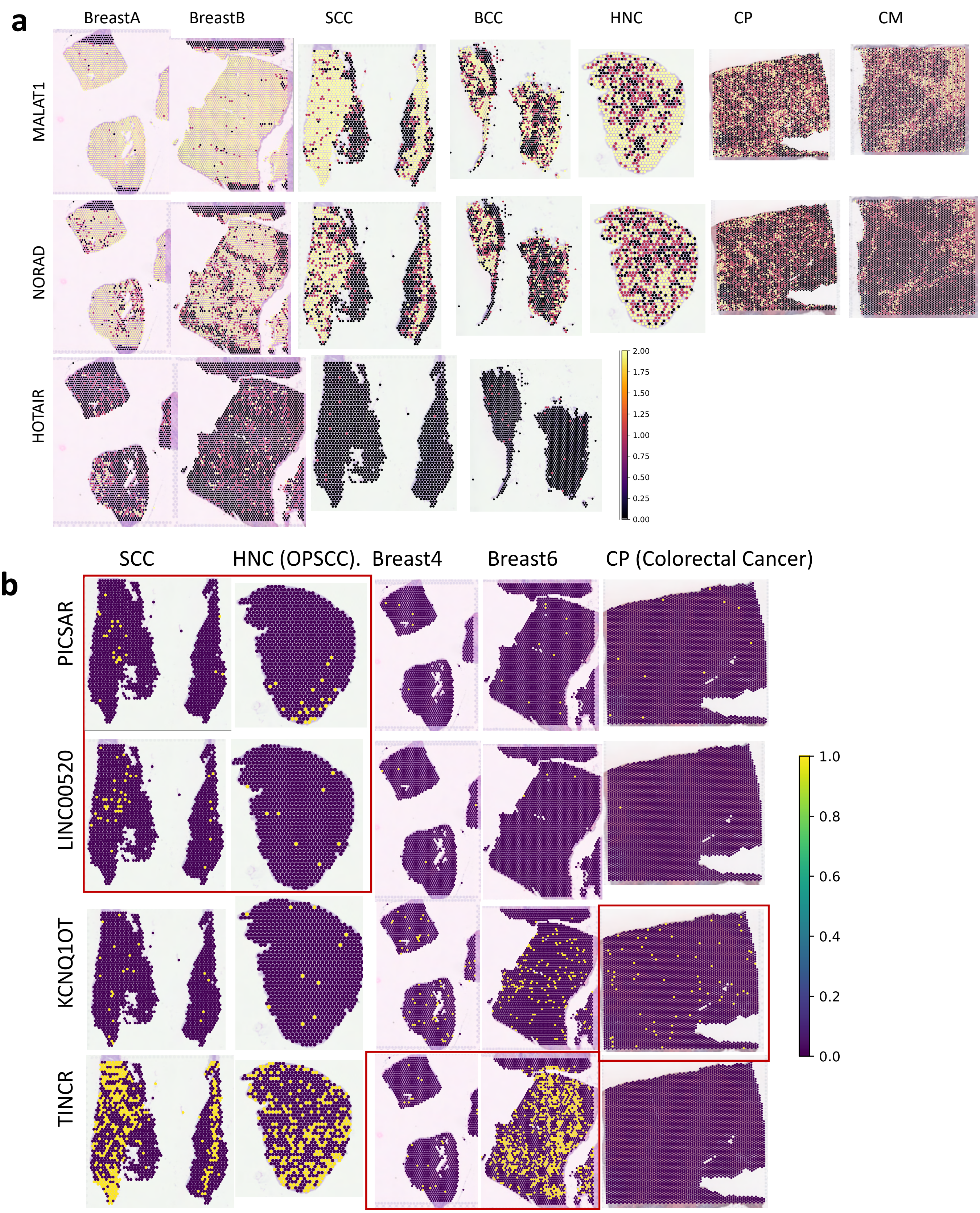


**Supplementary Fig. 3: a,** Expression of annotated well known lncRNAs in our in-house ST datasets **b,** Expression of annotated cancer-specific lncRNAs across respective cancer types in our in-house ST datasets

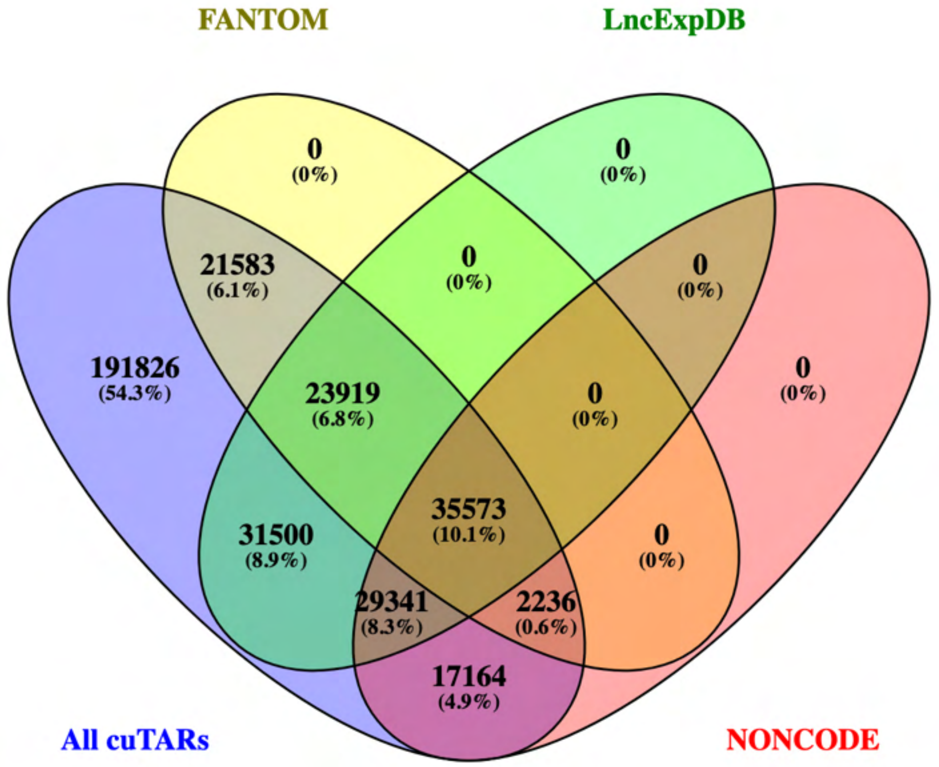


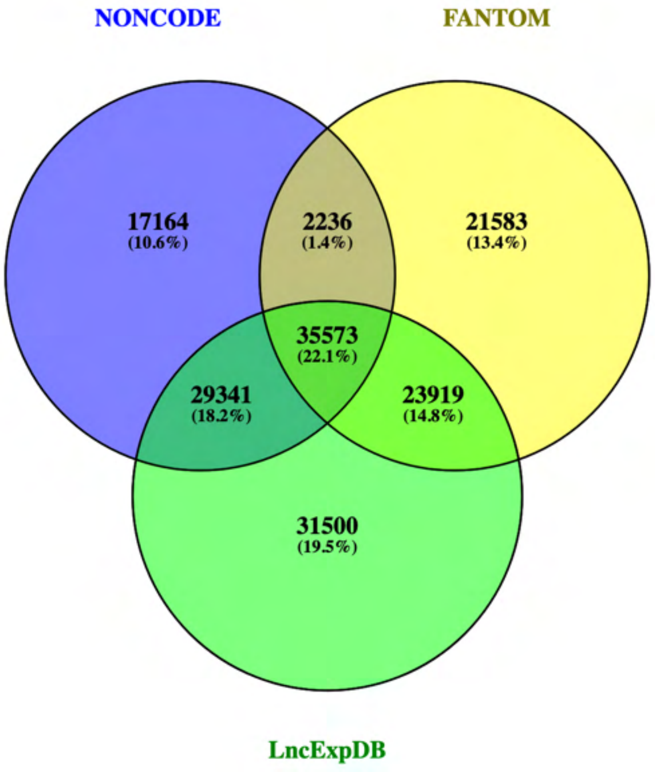


**Supplementary Fig. 4 :** cuTARs overlapping with NONCODE v6 lncRNAs

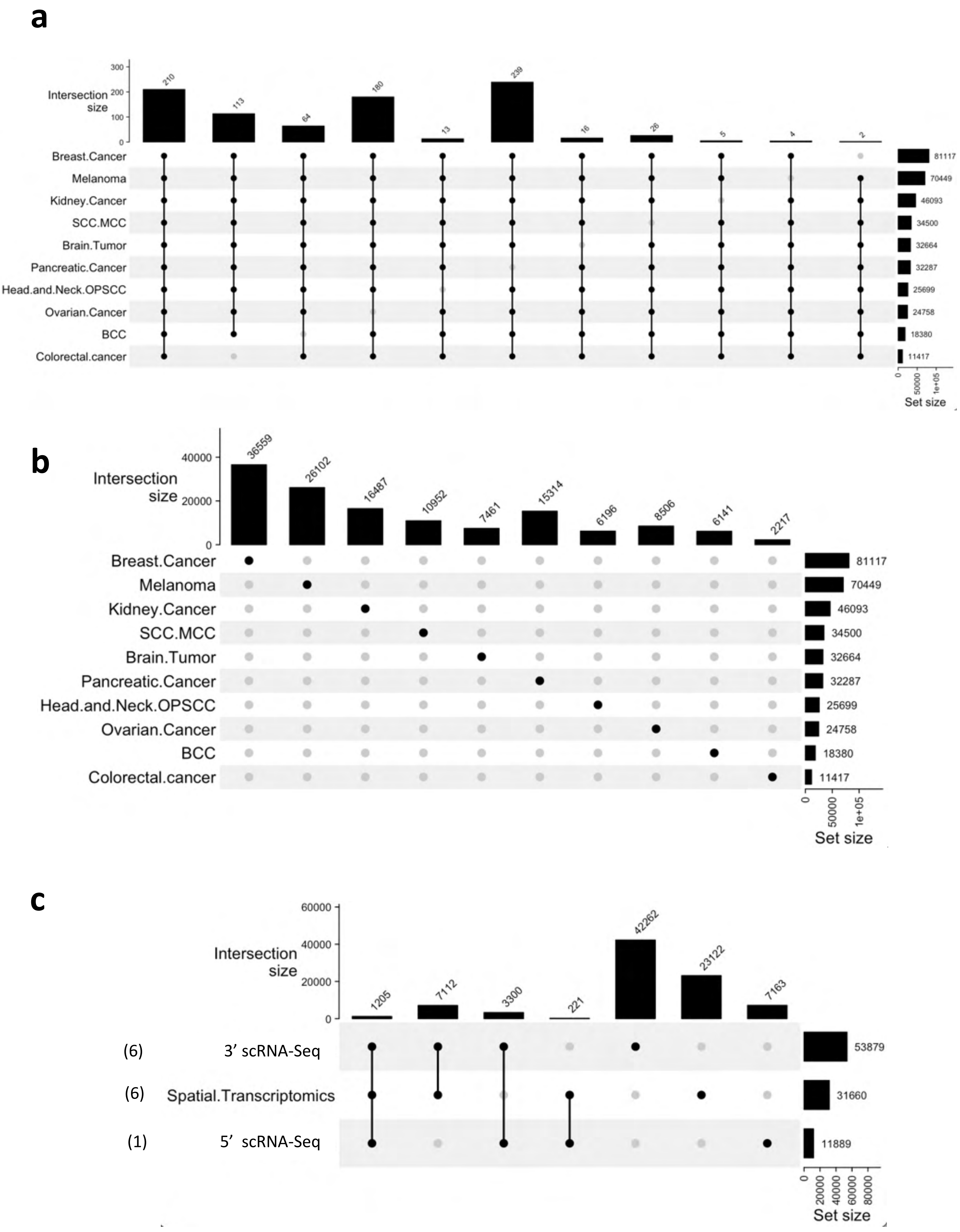


**Supplementary Fig. 5:** a, Pan-cancer cuTARs. **b,** cuTARs specific to each cancer type. **c,** Consistency of uTAR detection for different Melanoma samples across the three different sequencing platforms

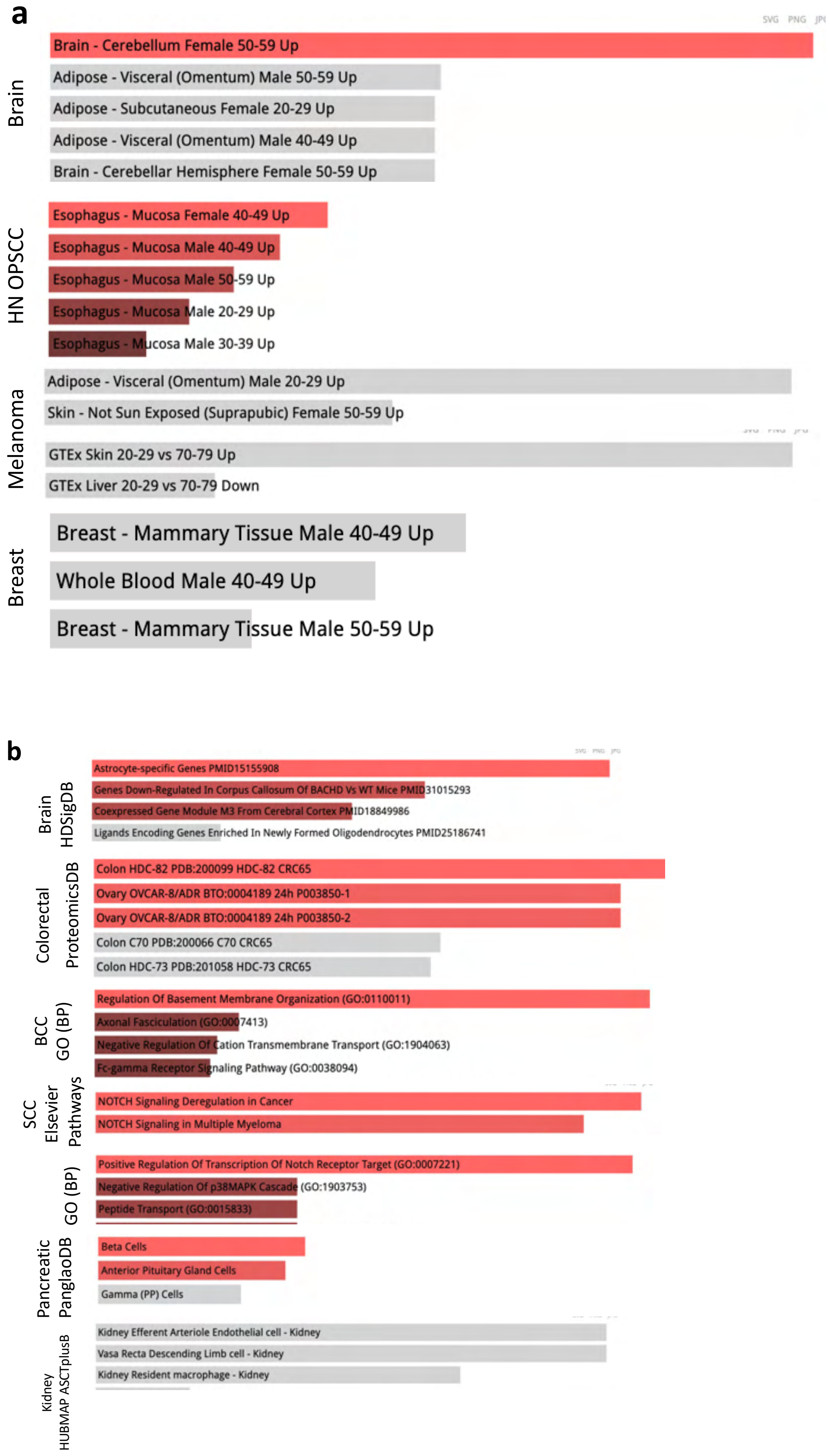


**Supplementary Fig. 6: a,** GTEx tissues enriched for genes associated with eQTLs colocalized with cancer-specific cuTARs. **b,** Other terms including pathways, GO, cell types

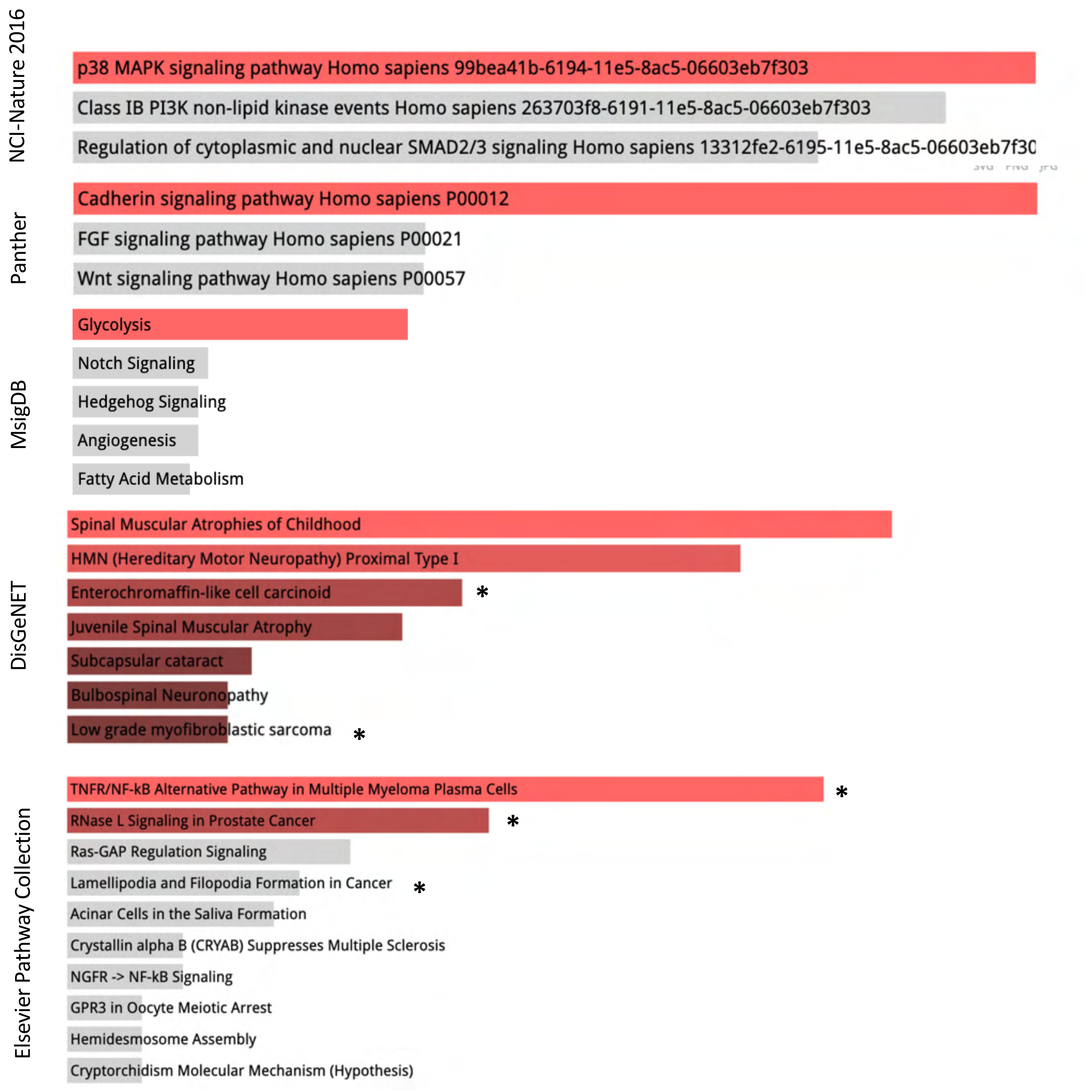


**Supplementary Fig. 7:** Pathways enriched for genes associated with eQTLs colocalized with pan-cancer cuTARs

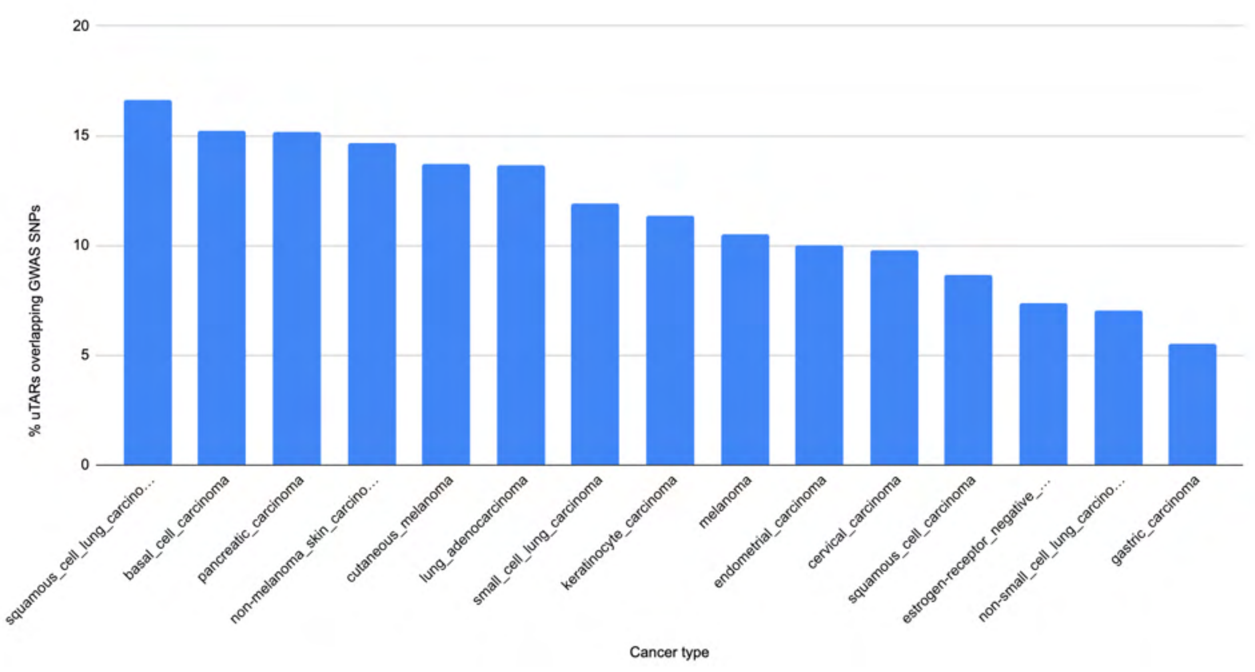


**Supplementary Fig. 8:** % uTARs from the in-house ST datasets overlapping with GWAS SNPs associated with different cancer types

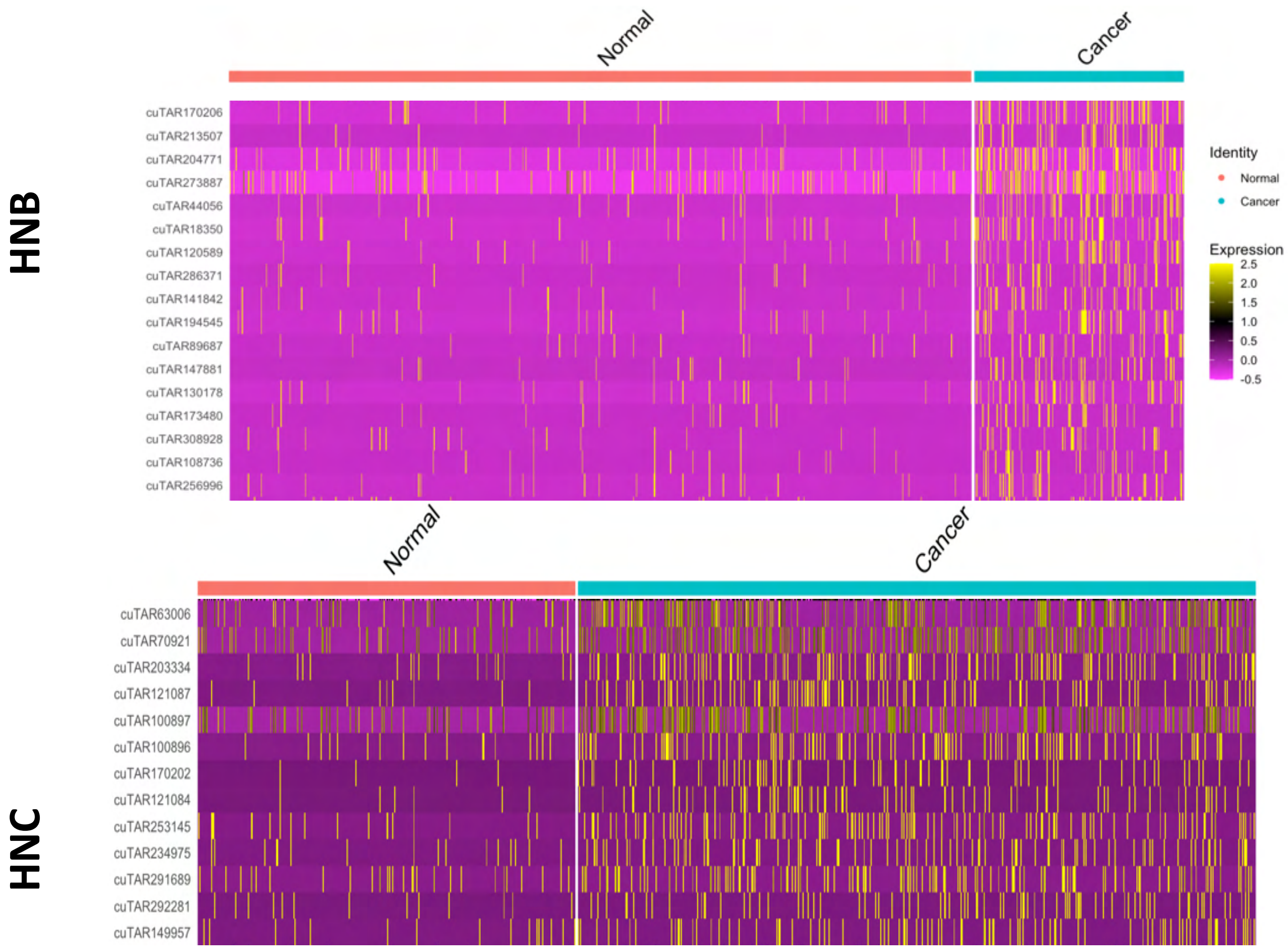


**Supplementary Fig. 9:** Differential uTAR expression in Head and Neck OPSCC samples

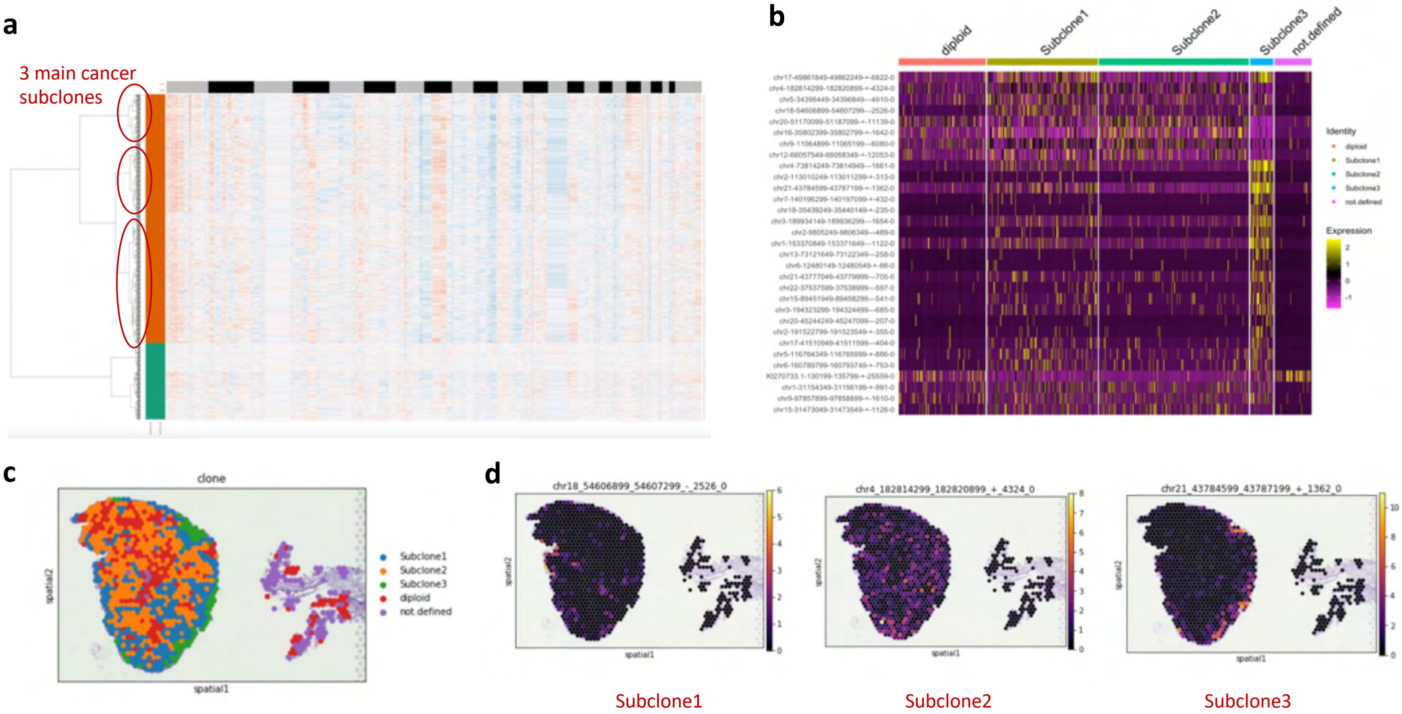


**Supplementary Fig. 10: a,**Three major cancer subclones identified using copyKAT. **b,** DE uTARs across subclones. **c,** Subclones annotated on the Head and Neck OPSCC tissue. **d,**Expression of Subclone-specific cuTARs

| Features | 10X Visium | No.of Features detected with ONT | %Features detected with ONT |
|---|---|---|---|
| **uTARs_uniq** | 3025 | 834 + 166 | 27.5702 |
| **Genes** | 16523 | 12336 | 74.6595 |

**Supplementary Table 1**: 10X Visium *versus* Nanopore stats

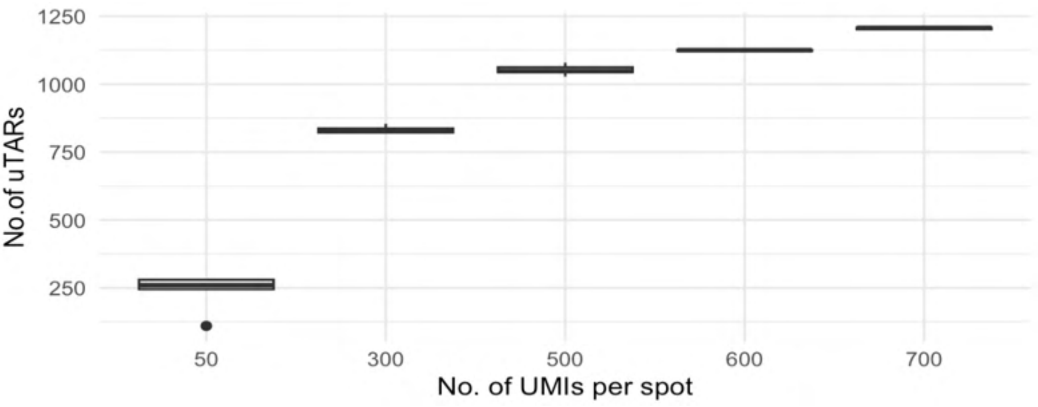


b

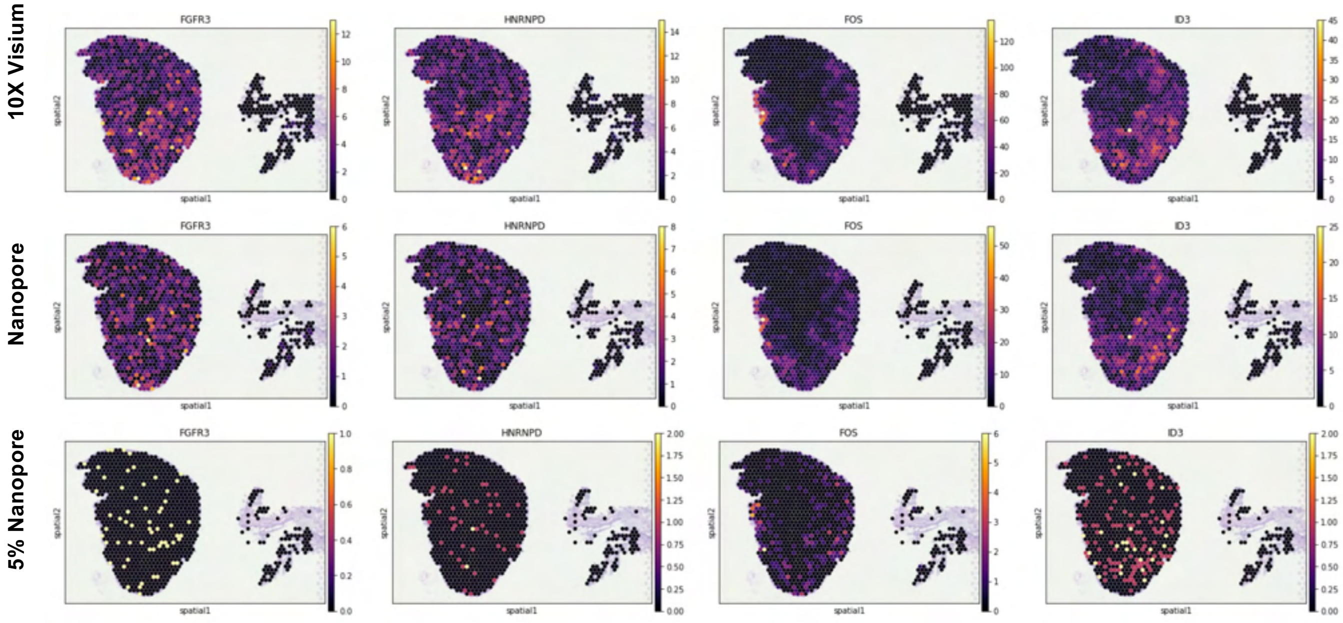


**Supplementary Fig. 11 : a,** No. of UTARs for downsampled number of unique molecules per spot. **b,** Gene expression detected from 10X Visium and ONT

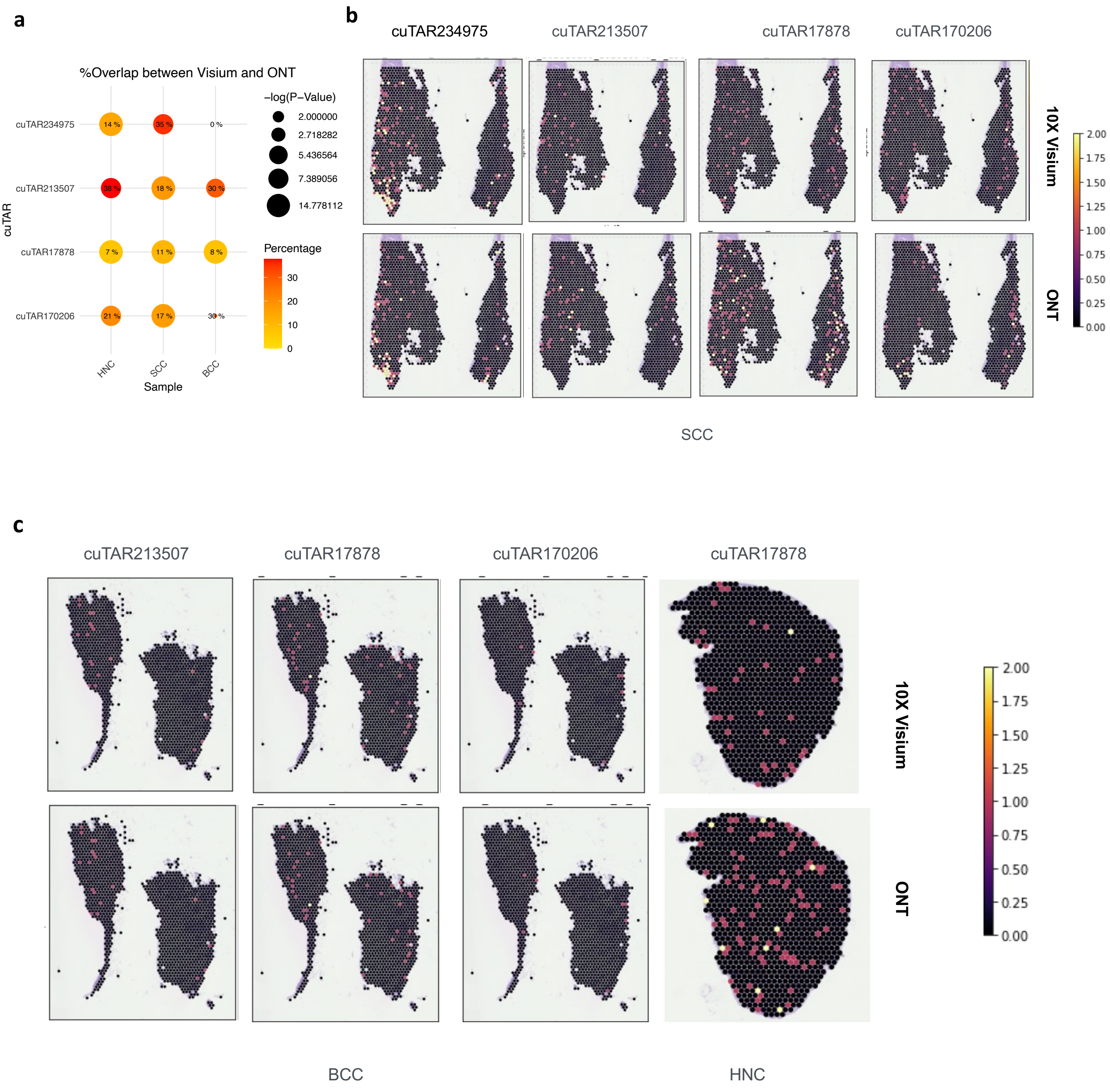


**Supplementary Fig. 12 : a,** % Positive spots expressing PCR validated cuTARs across Visium and ONT. **b,** Spatial plots for these cuTARs in SCC across the two platforms **c,** Spatial plots for these cuTARs in BCC and HNC across the two platforms

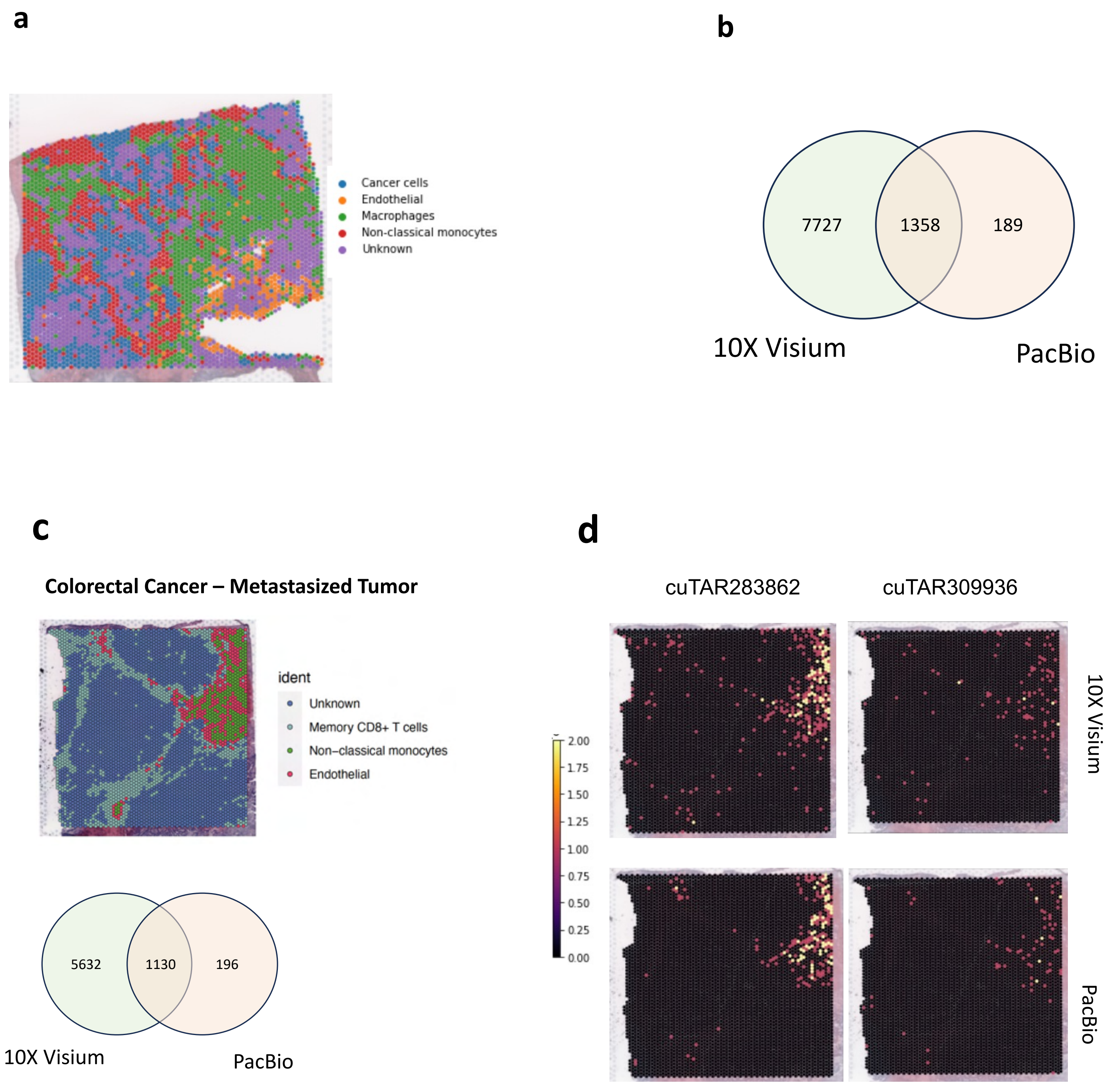


**Supplementary Fig. 13** : **a,** Cell types in Primary colorectal cancer tissue. **b,** uTARs validated with long read HiFi sequencing by PacBio **c,** Cell types in Metastasized colorectal cancer tissue and number uTARs validated with long read HiFi sequencing by PacBio. **d,** Expression of two cuTARs across both platforms

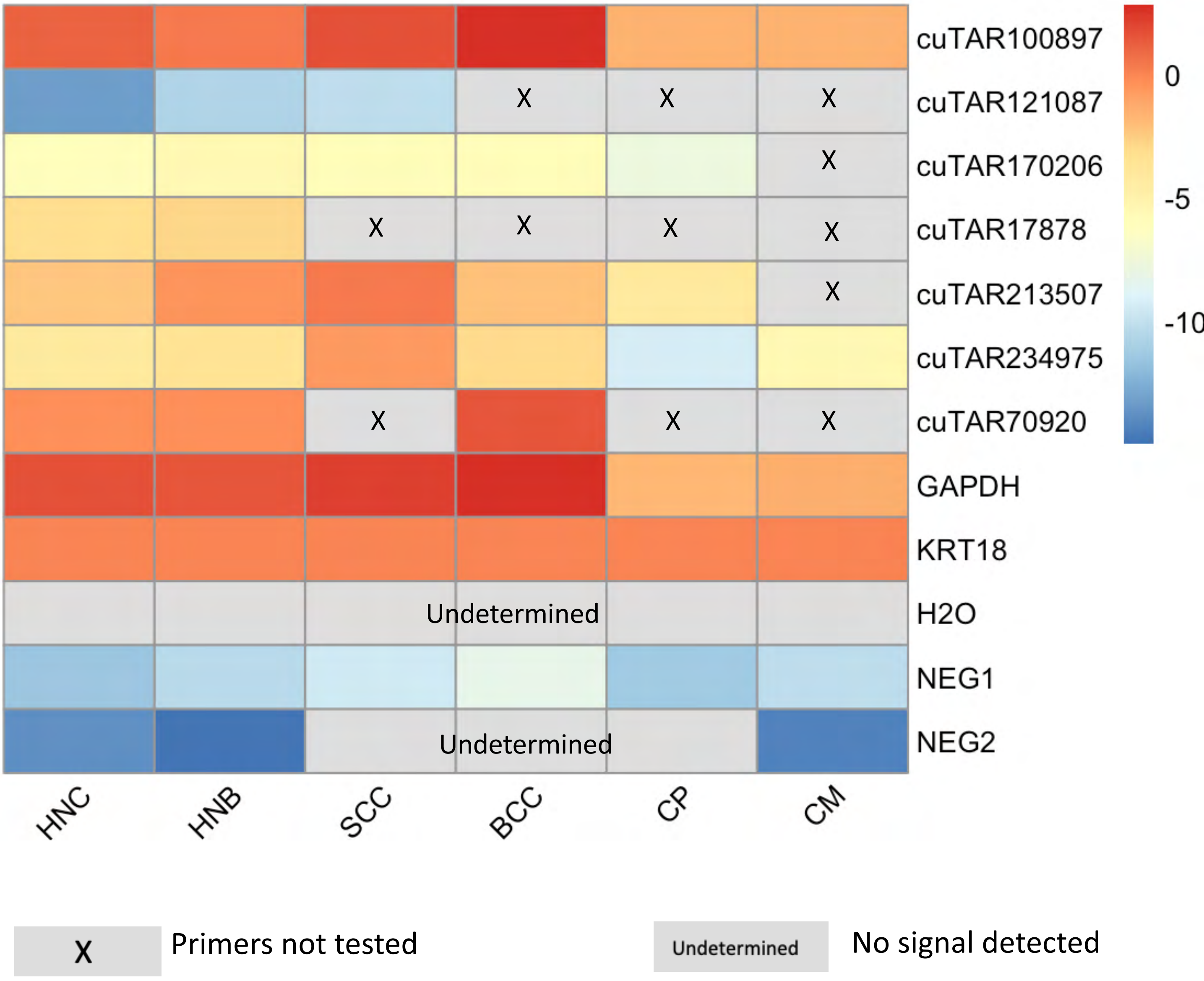


**Supplementary Fig. 14:** Expression of some cuTARs across samples using qPCR

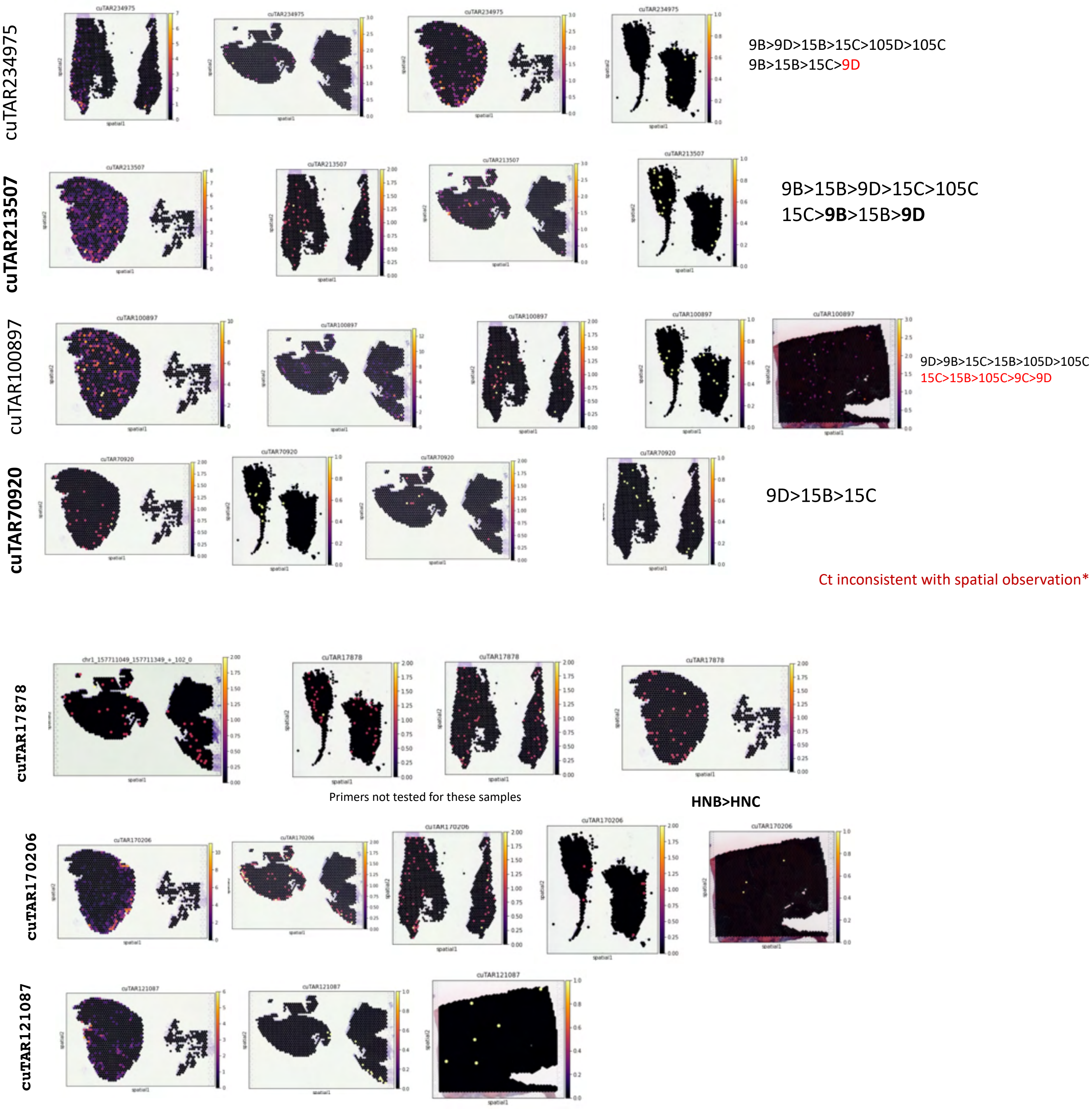


**Supplementary Fig. 15:** 10X Visium Expression of the qPCR validated cuTARs

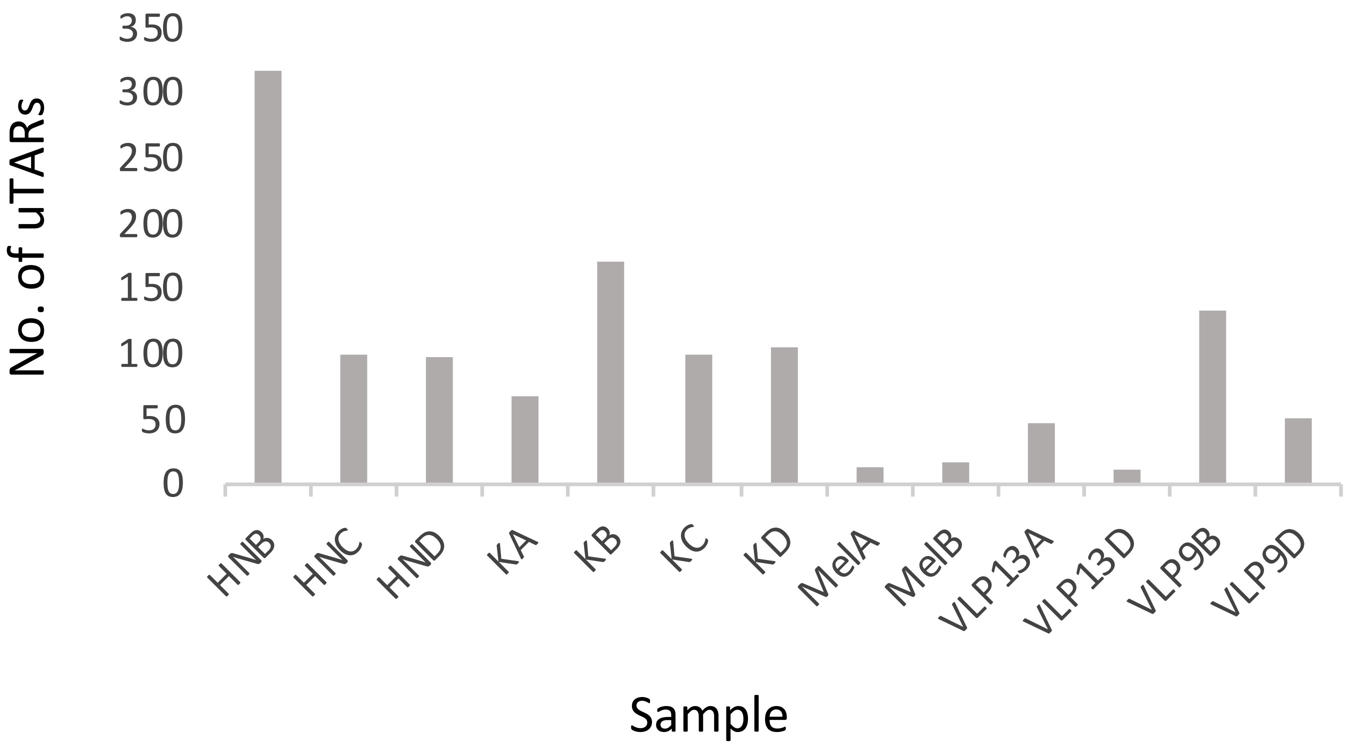


**Supplementary Fig. 16** : Number of spatially variable uTARs identified using SpaDE

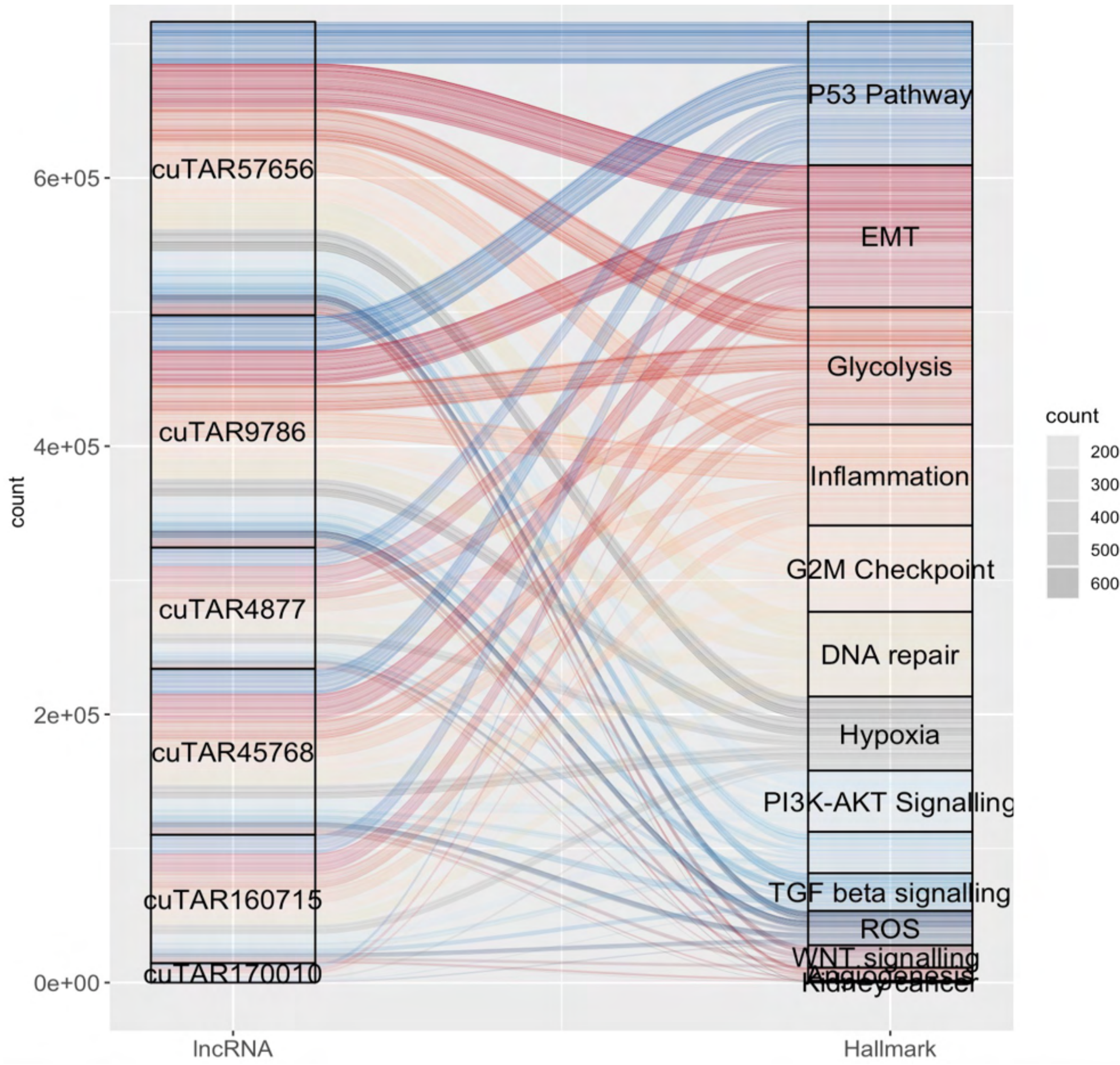


**Supplementary Fig. 17:** Top uTARs in kidney cancer (sample A) with high spatial autocorrelation with different cancer-relevant gene sets

**a**

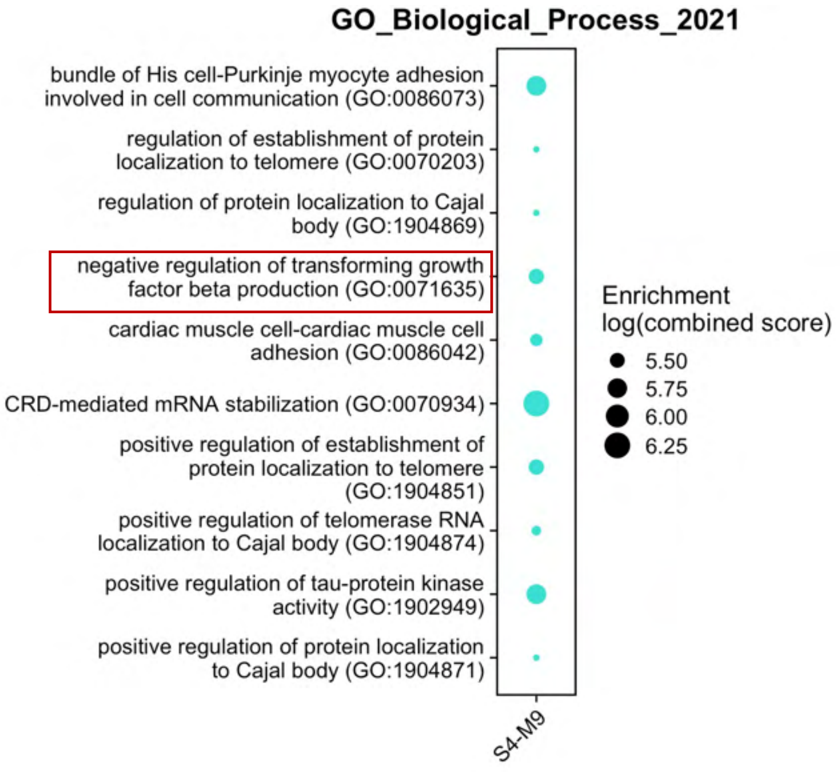


**b**

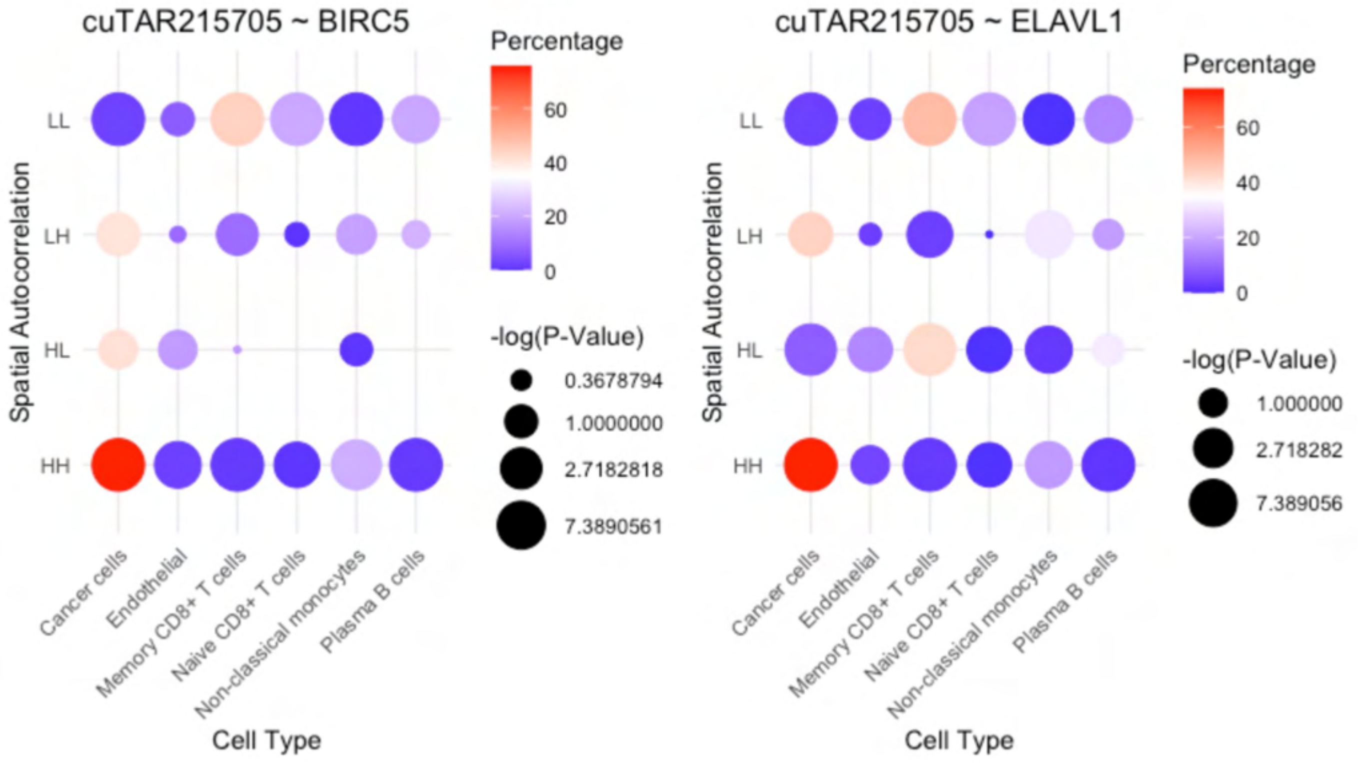


**Supplementary Fig. 18: a,** GO Terms associated with a co-expression module with cuTAR215705. **b,** Spatial autocorrelation of the cuTAR with interacting gene and RBP across cell-types

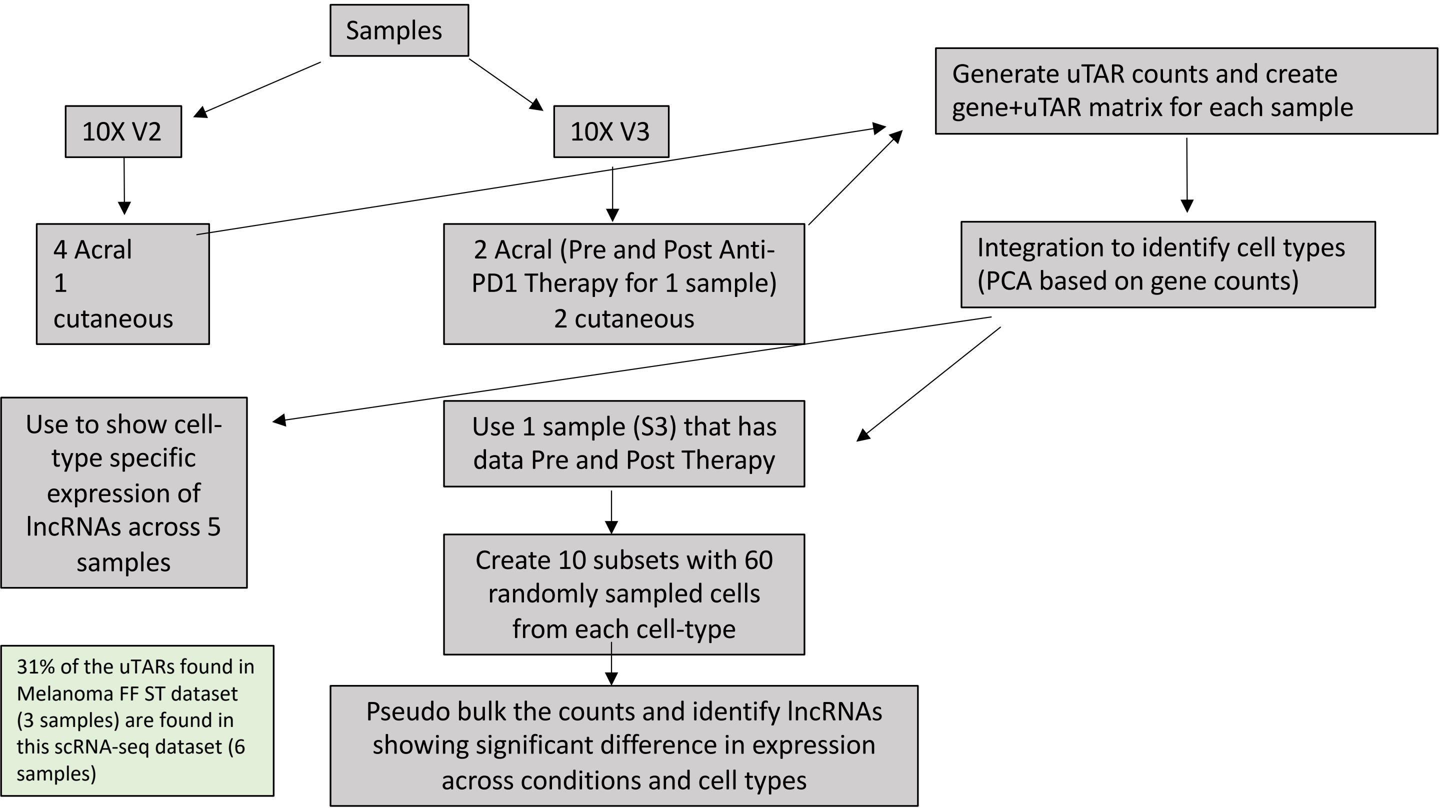


**Supplementary Fig. 19:** Single-cell Melanoma data analysis workflow

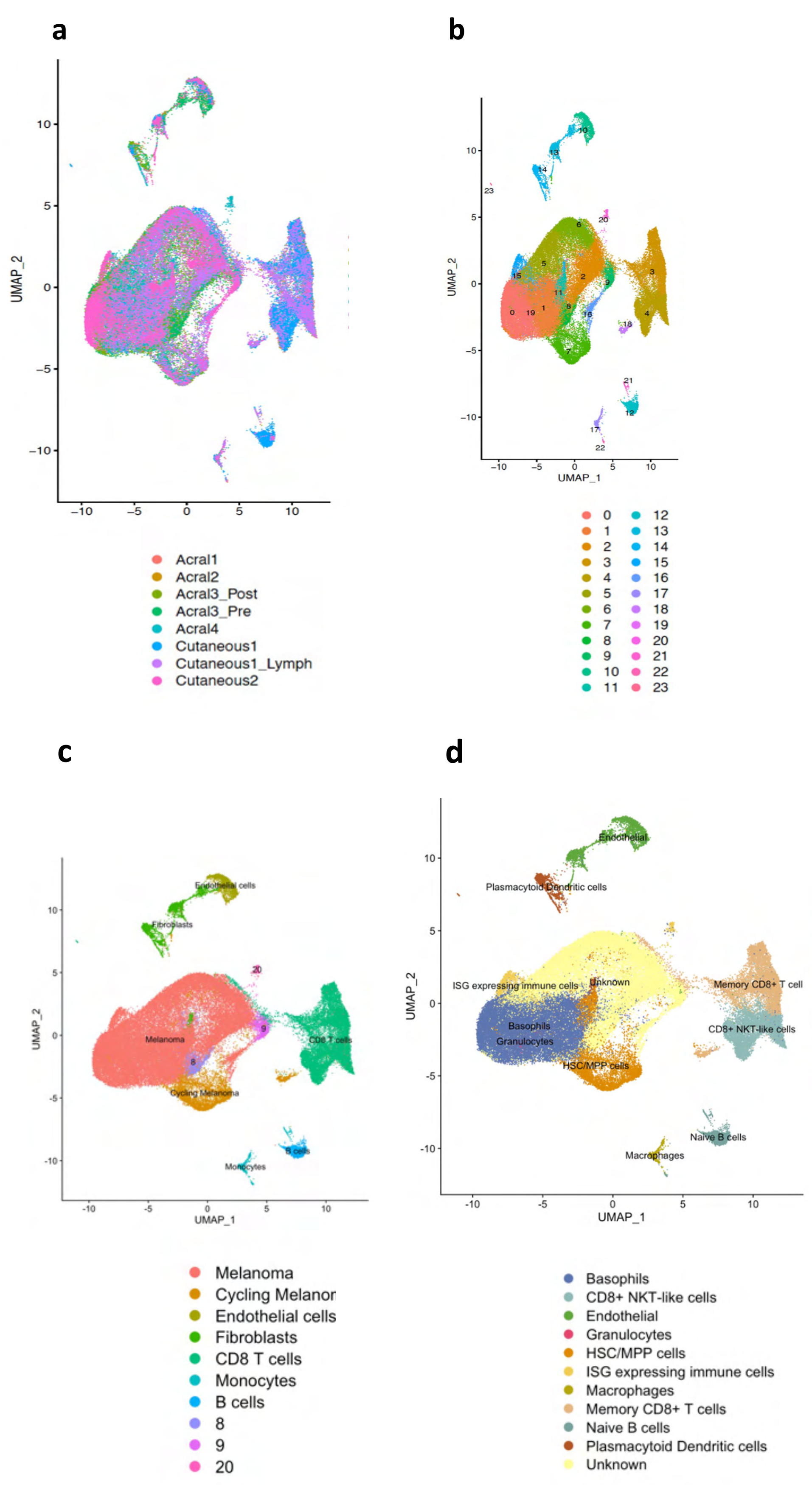


**Supplementary Fig. 20: a,** All samples from the study integrated. **b,** Seurat clusters. **c,** Manual annotation based on Marker genes from the paper. **d,** Automated annotation with scType

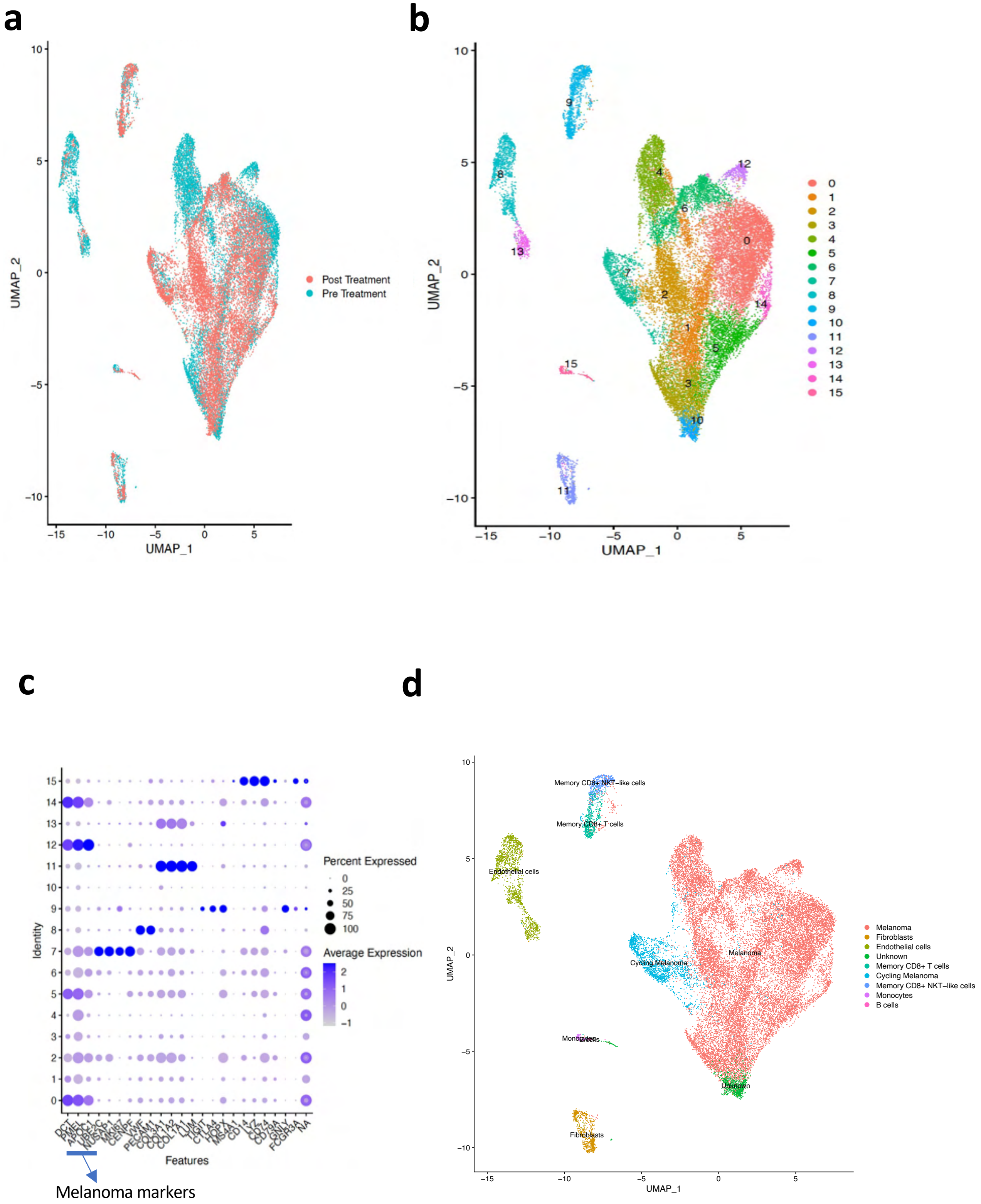


**Supplementary Fig. 21: Analysis of an Acral melanoma sample – Pre and post treatment a,** Integration of 2 samples (S3 Pre and Post). **b,** Clustering. **c,** Canonical cell-type marker expression. **d,** cluster annotations

**a**

**b**

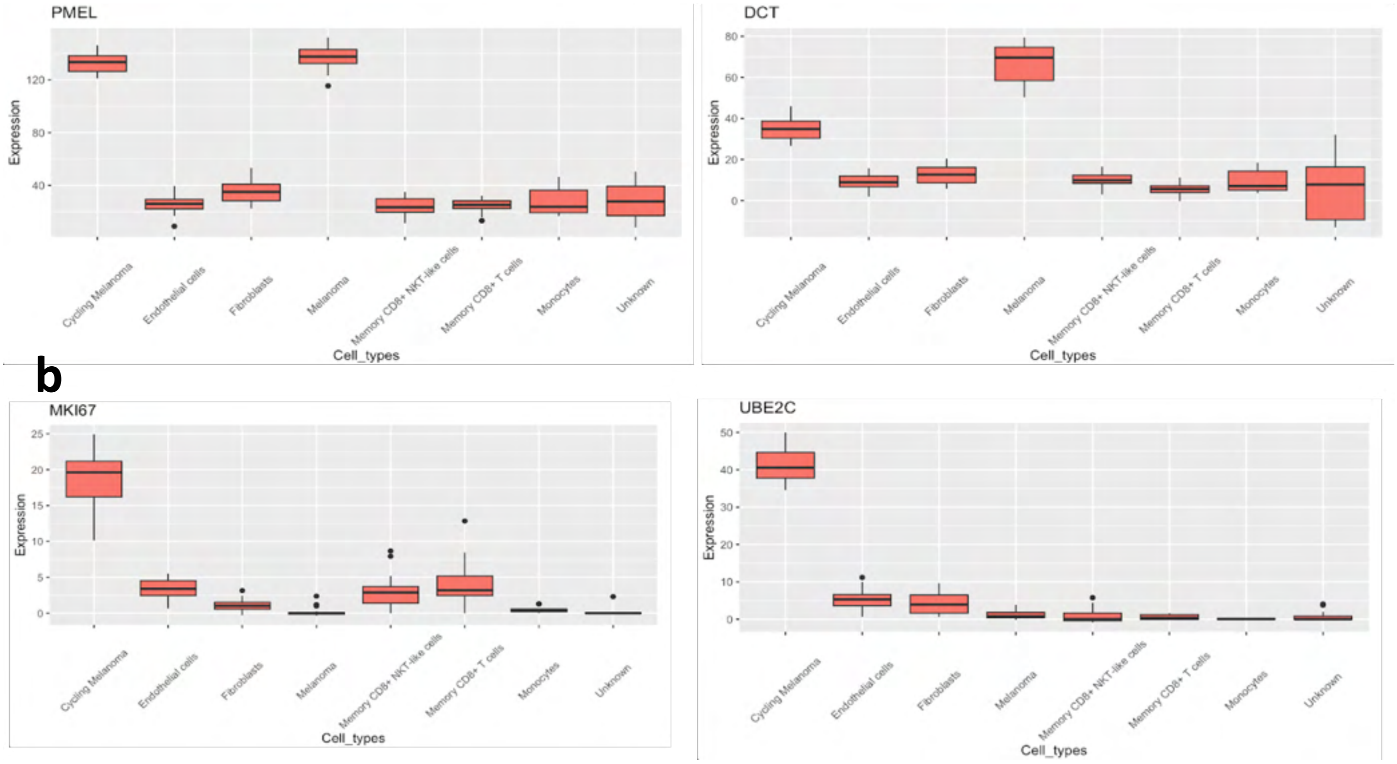


**Supplementary Fig. 22:** Expression of **a,** Melanoma specific genes and **b,** proliferation markers across various cell types.

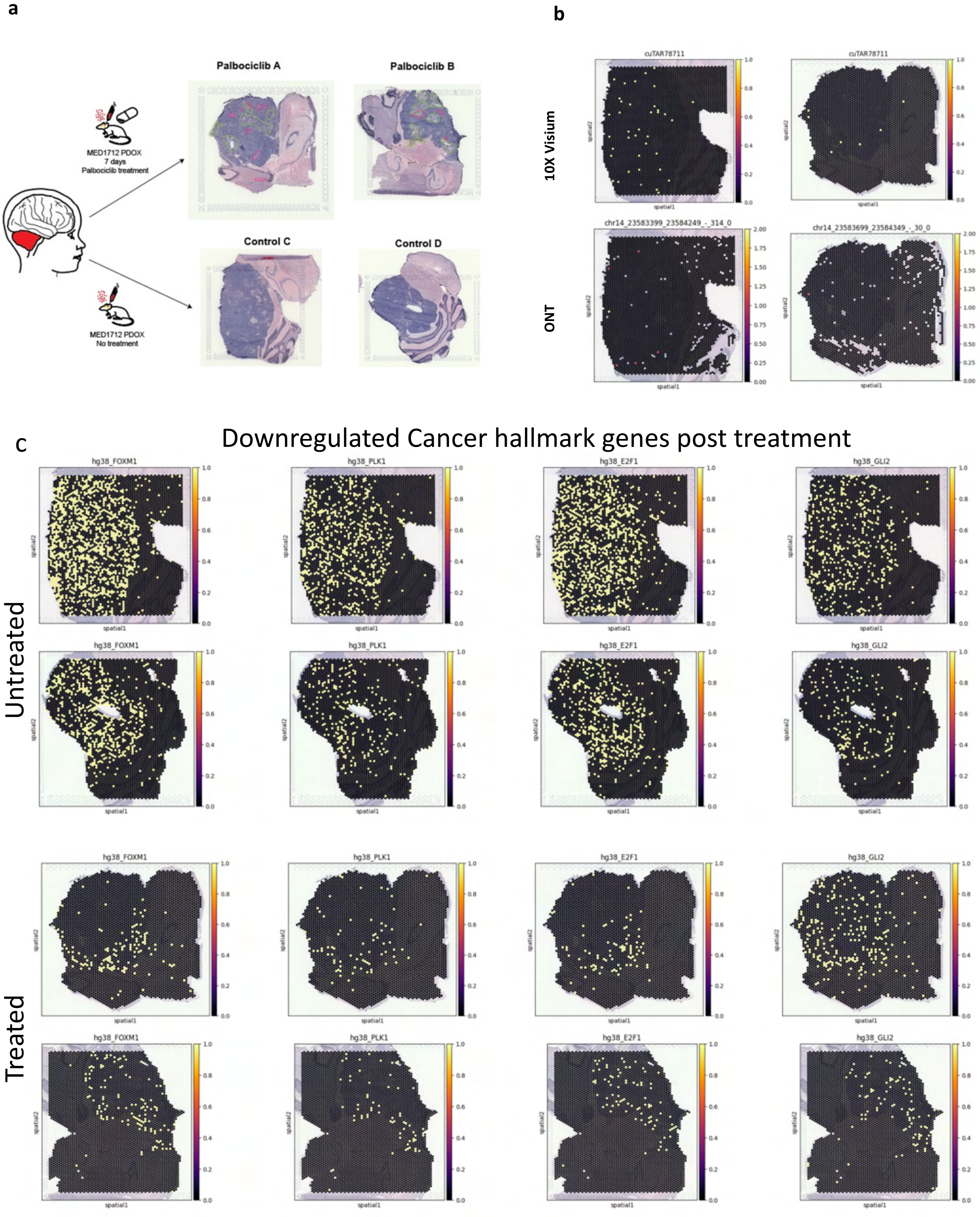


**Supplementary Fig. 23: a,** Tissues from PDOX mouse models. **b,** Expression of the cuTARs across Visium and ONT **c,** Expression of cancer hallmark genes in untreated and treated samples

### cuTAR67350

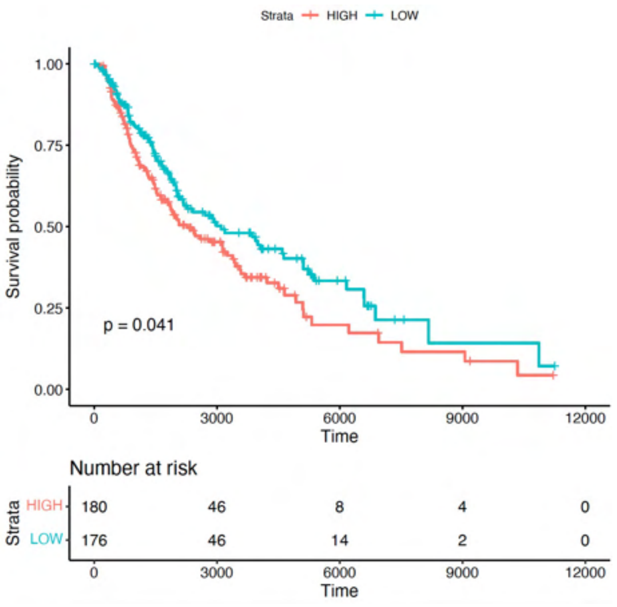


### cuTAR293520

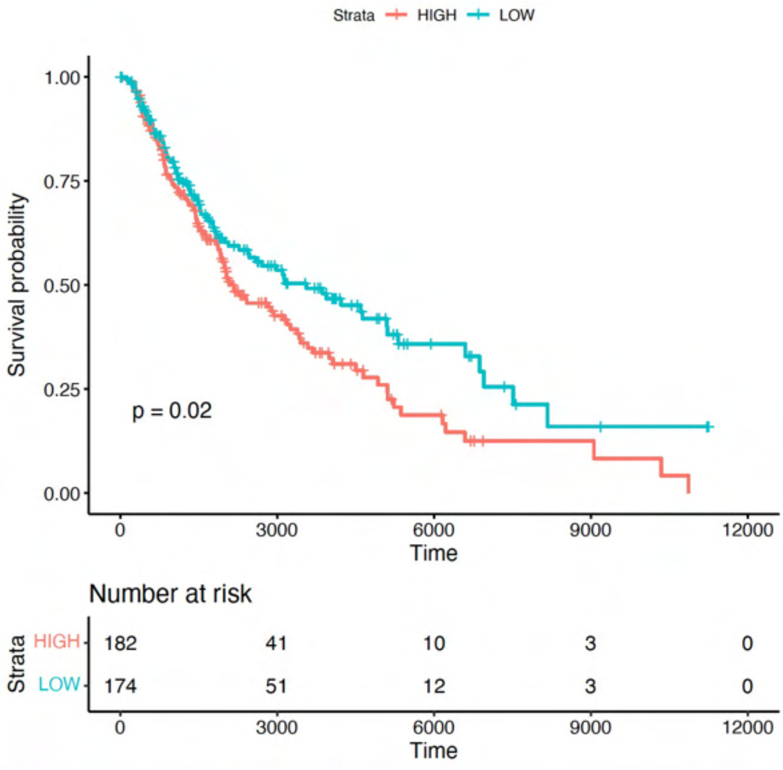


**Supplementary Fig. 24:** Kaplan-Meyer curves for cuTARs showing significant differences in survival probabilities for high and low expressing groups

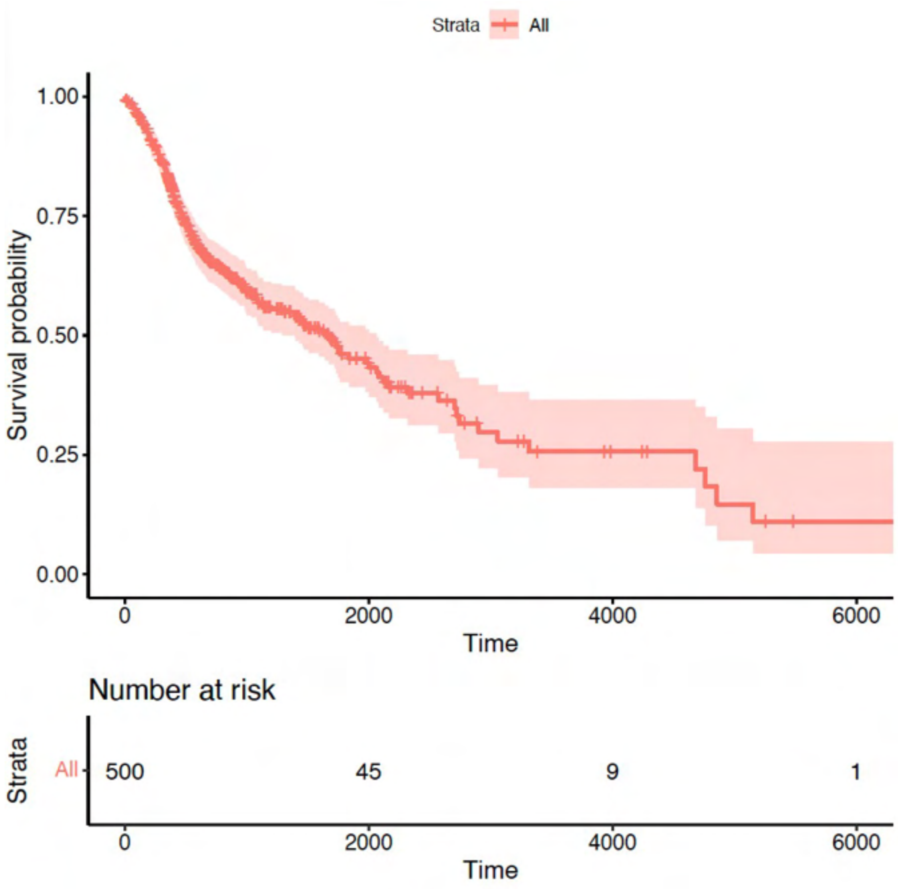


**Supplementary Fig. 25**: No stratification in Head and Neck cancer samples for the Melanoma-specific cuTARs. All samples were low expressing

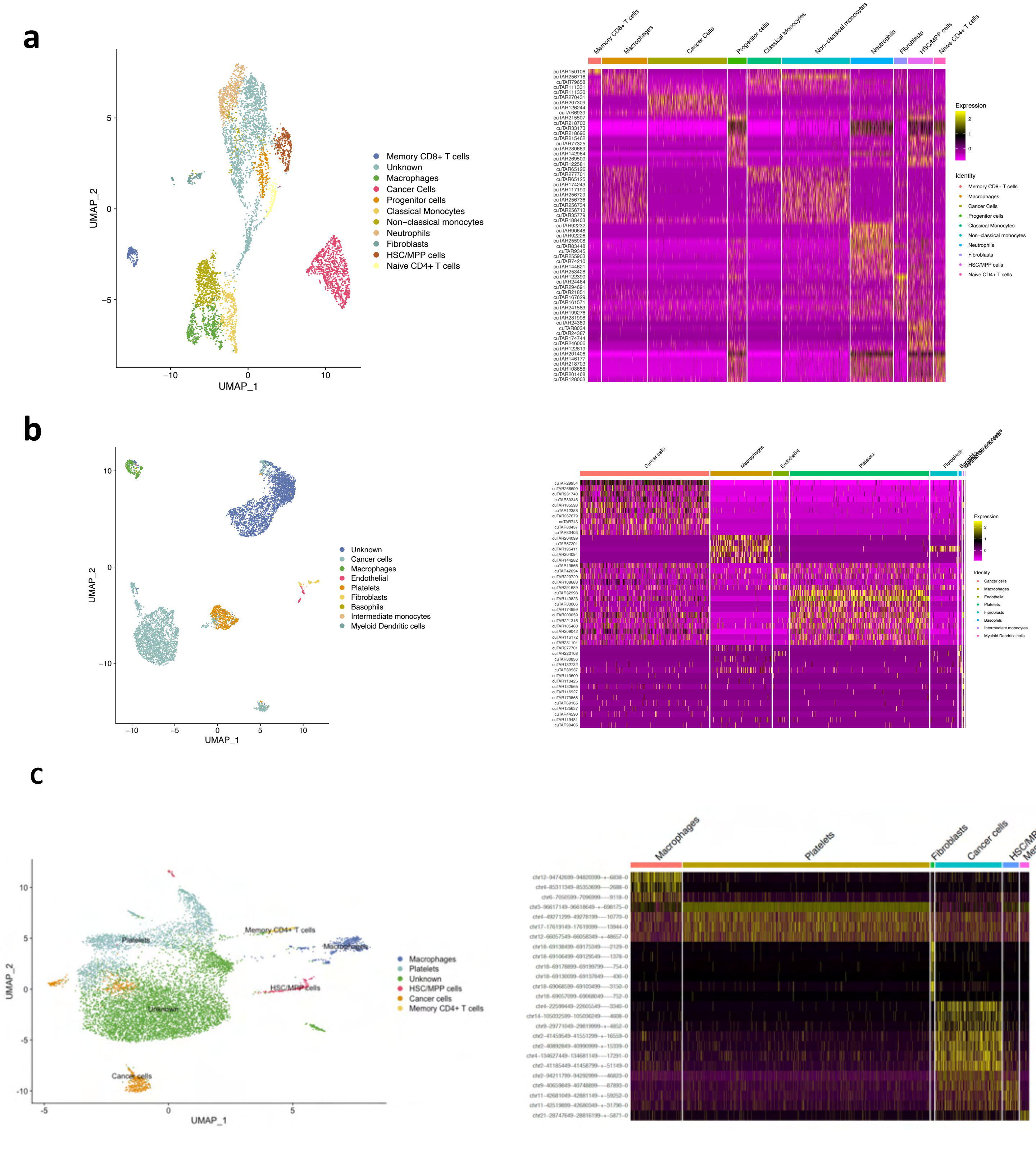


**Supplementary Fig. 26:** Clustering and cell-type specific cuTARs in **a,** Glioblastoma **b,** Breast cancer (Invasive Ductal carcinoma) **c**, Kidney cancer

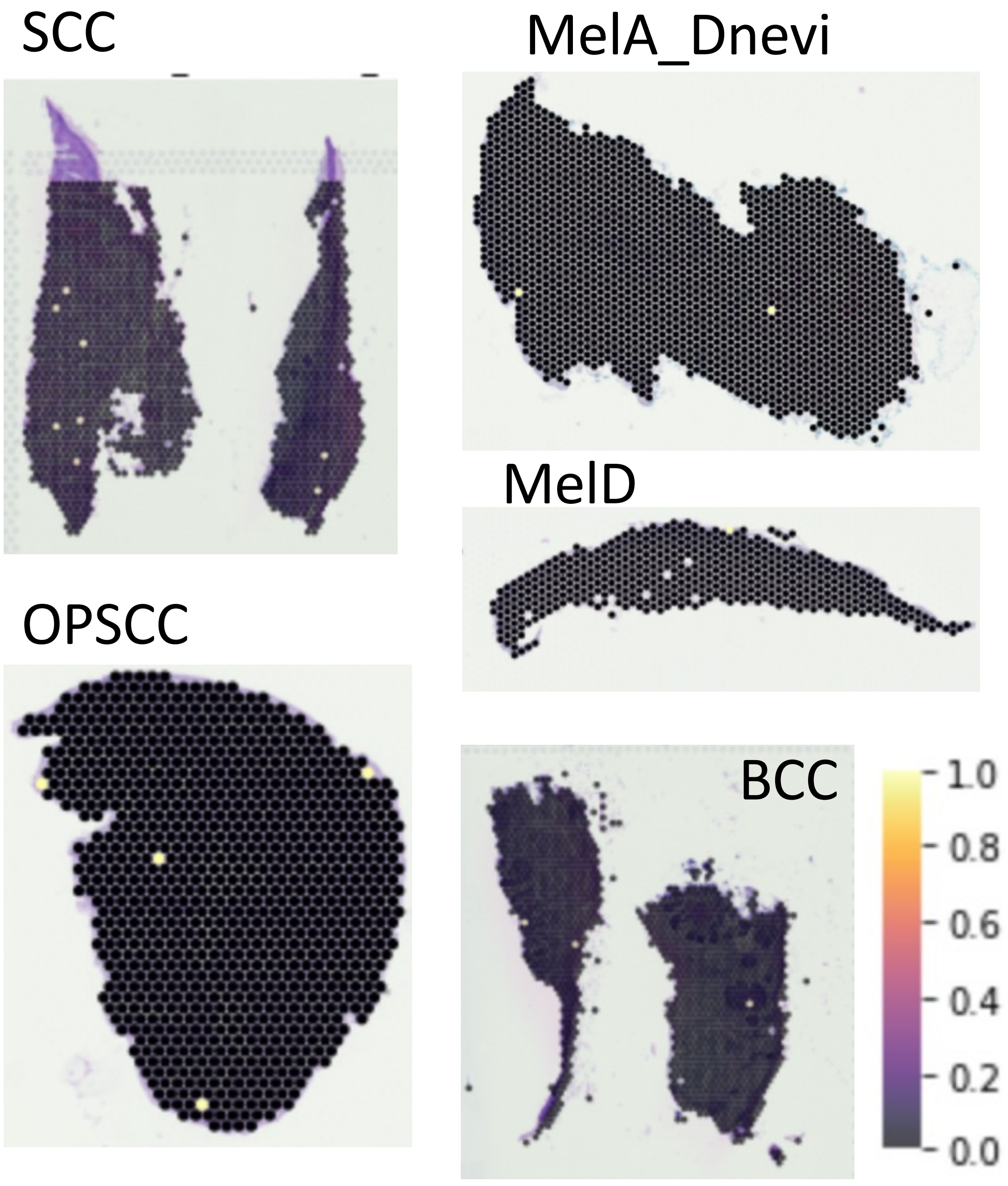


**Supplementary Fig. 27:** cuTAR87324, upregulated in the tumor cells of kidney cancer is detected across OPSCC (validated by ONT) and skin cancers although very low expression is seen
